# An instantaneous voice synthesis neuroprosthesis

**DOI:** 10.1101/2024.08.14.607690

**Authors:** Maitreyee Wairagkar, Nicholas S. Card, Tyler Singer-Clark, Xianda Hou, Carrina Iacobacci, Leigh R. Hochberg, David M. Brandman, Sergey D. Stavisky

**Affiliations:** Department of Neurological Surgery, University of California Davis, Davis, CA; Department of Biomedical Engineering, University of California Davis, Davis, CA; Department of Computer Science, University of California Davis, Davis, CA; School of Engineering and Carney Institute for Brain Sciences, Brown University, Providence, RI; VA RR&D Center for Neurorestoration and Neurotechnology, VA Providence Healthcare, Providence, RI; Center for Neurotechnology and Neurorecovery, Department of Neurology, Massachusetts General Hospital, Harvard Medical School, Boston, MA

## Abstract

Brain computer interfaces (BCIs) have the potential to restore communication to people who have lost the ability to speak due to neurological disease or injury. BCIs have been used to translate the neural correlates of attempted speech into text^1–3^. However, text communication fails to capture the nuances of human speech such as prosody, intonation and immediately hearing one’s own voice. Here, we demonstrate a “brain-to-voice” neuroprosthesis that instantaneously synthesizes voice with closed-loop audio feedback by decoding neural activity from 256 microelectrodes implanted into the ventral precentral gyrus of a man with amyotrophic lateral sclerosis and severe dysarthria. We overcame the challenge of lacking ground-truth speech for training the neural decoder and were able to accurately synthesize his voice. Along with phonemic content, we were also able to decode paralinguistic features from intracortical activity, enabling the participant to modulate his BCI-synthesized voice in real-time to change intonation, emphasize words, and sing short melodies. These results demonstrate the feasibility of enabling people with paralysis to speak intelligibly and expressively through a BCI.

## Introduction

Speaking is an essential human ability, and losing the ability to speak is devastating for people living with neurological disease and injury. Brain computer interfaces (BCIs) are a promising therapy to restore speech by bypassing the damaged parts of the nervous system through decoding neural activity^4^. Recent demonstrations of BCIs have focused on decoding neural activity into text on a screen^2,3^ with high accuracy^1^. While these approaches offer an intermediate solution to restore communication, communication with text alone falls short of providing a digital surrogate vocal apparatus with closed-loop audio feedback and fails to restore critical nuances of human speech including prosody.

These additional capabilities can be restored with a “brain-to-voice” BCI that decodes neural activity into sounds in real-time that the user can hear as they attempt to speak. Developing such a speech synthesis BCI poses several unsolved challenges: the lack of ground truth training data, i.e., not knowing how and when a person with speech impairment is trying to speak; causal low-latency decoding for instantaneous voice synthesis that provides continuous closed-loop audio feedback; and a flexible decoder framework for producing unrestricted vocalizations and modulating paralinguistic features of the synthesized voice.

A growing literature of studies have reconstructed voice offline from able-bodied speakers using previously recorded neural signals measured with electrocorticography (ECoG)^5–11^, stereoencephalography (sEEG)^12^, and intracortical microelectrode arrays^13,14^. Decoders trained on overt speech of able speakers could synthesize unintelligible speech during miming, whispering or imaging speaking tasks online^8,15^ and offline^16^. Recently, intermittently intelligible speech was synthesized seconds after a user with ALS spoke overtly (and intelligibly)^17^ from a six-word vocabulary. While the aforementioned studies were done with able speakers, a study ^3^ with a participant with anarthria adapted a text decoding approach to decode discrete speech units acausally at the end of the sentence to synthesize speech from a 1,024-word vocabulary. However, this is still very different from healthy speech, where people immediately hear what they are saying and can use this to accomplish communication goals such as interjecting in a conversation. In this work, we sought to synthesize voice continuously and with low latency from neural activity as the user attempted to speak, which we refer to as “instantaneous” voice synthesis to contrast it with earlier work demonstrating acausal delayed synthesis.

Here, we report an instantaneous brain-to-voice BCI using 256 microelectrodes chronically placed in the precentral gyrus of a man with severe dysarthria due to amyotrophic lateral sclerosis (ALS). We did not have ground-truth voice data from this participant. To overcome this, we generated synthetic target speech waveforms from the prompt text and time-aligned these with neural activity to estimate the participant’s intended speech. We were then able to train a deep learning model that synthesized his intended voice in real-time by decoding his neural activity causally within 10 ms. The resulting synthesized voice was often (but not consistently) intelligible and human listeners were able to identify the words with high accuracy. This flexible brain-to-voice framework – which maps neural activity to acoustic features without an intermediary such as discrete speech tokens or limited vocabulary – could convert participant’s neural activity to a realistic representation of his pre-ALS voice, demonstrating voice personalization, and it enabled the participant to speak out-of-dictionary pseudo-words and make interjections.

We also found that in addition to previously-documented phonemic information^1,2^, there is substantial paralinguistic information in the intracortical signals recorded from ventral precentral gyrus. These features were causally decoded to enable the participant to modulate his BCI voice to change intonation in order to ask a question or emphasize specific words in a sentence, and to sing melodies with different pitch targets. Finally, we investigated the dynamics of the neural ensemble activity, which revealed that putatively output-null neural dimensions are highly active well before each word is vocalized, with greater output-null activity present when there were more upcoming words planned and when the upcoming word needed to be modulated.

## Results

### Continuous speech synthesis from intracortical neural activity with immediate auditory feedback

We recorded neural activity from four microelectrode arrays with a total of 256 electrodes placed in the ventral premotor cortex (6v), primary motor cortex (M1) and middle precentral gyrus (55b) (see **Fig. 1a,b**) as estimated using the Human Connectome Project pipeline^1,18^ in BrainGate2 clinical trial participant ‘T15’ (**Extended Data Fig. 1**). T15 was a 45-year-old man with ALS and severe dysarthria. He retained some orofacial movement and an ability to vocalize but was unable to produce intelligible speech (**Video 1**).

**Fig. 1.**
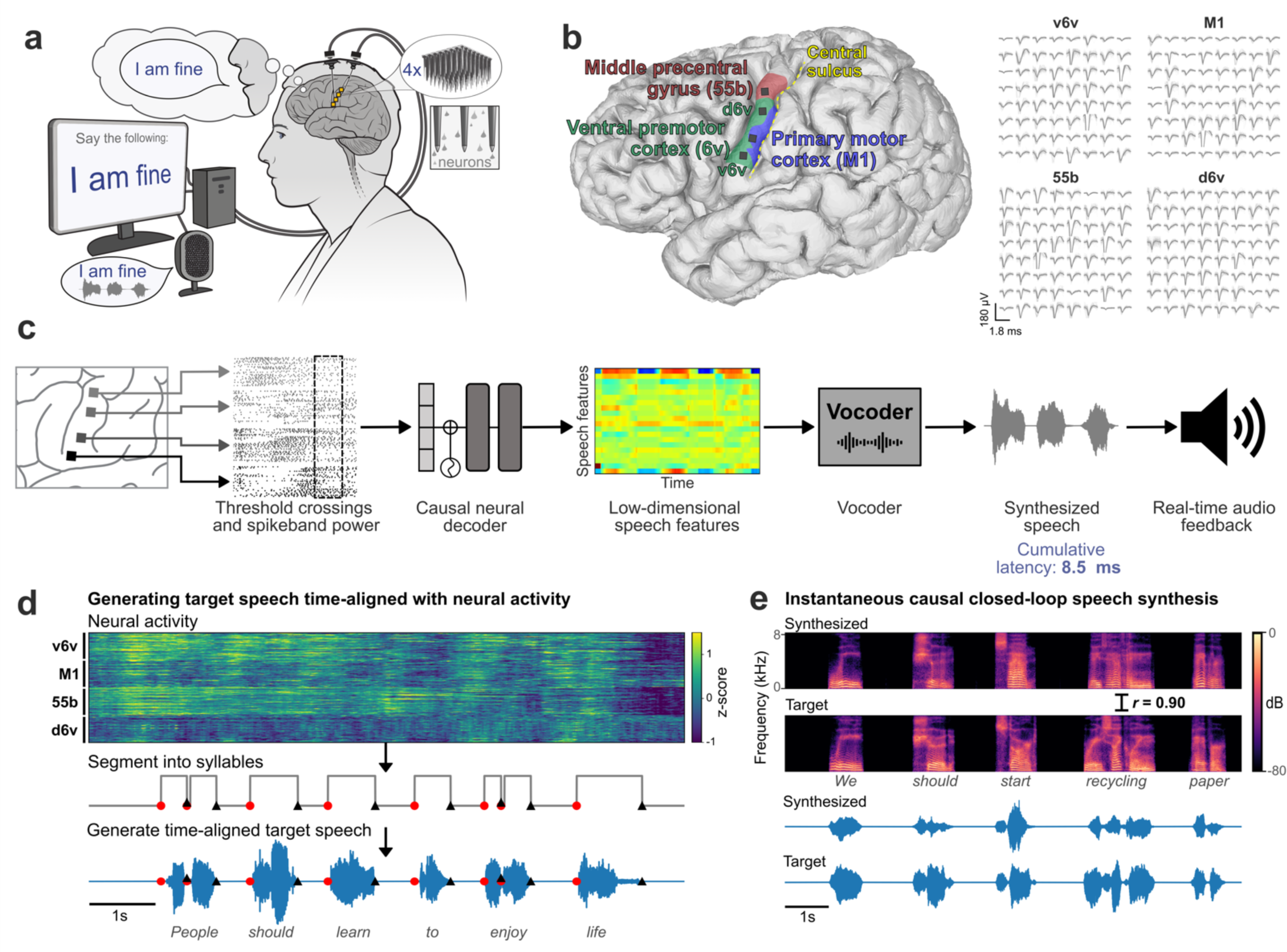
Closed-loop voice synthesis from intracortical neural activity in a participant with ALS. **a.** Schematic of the brain-to-voice neuroprosthesis. Neural features extracted from four chronically implanted microelectrode arrays were decoded in real-time and used to directly synthesize voice. **b.** Array locations on the participant’s left hemisphere and typical neuronal action potentials from each microelectrode. Color overlays are estimated from a Human Connectome Project cortical parcellation. **c.** Closed-loop causal voice synthesis pipeline: voltages were sampled at 30 kHz; threshold-crossings and spike-band power features were extracted from 1 ms segments; these features were binned into 10 ms non-overlapping bins, normalized and smoothed. The Transformer-based decoder mapped these neural features to a low-dimensional representation of speech involving Bark-frequency cepstral coefficients, pitch, and voicing, which were used as input to a vocoder. The vocoder then generated speech samples which were continuously played through a speaker. **d.** Lacking T15’s ground truth speech, we first generated synthetic speech from the known text cue in the training data using a text-to-speech algorithm, and then used the neural activity itself to time-align the synthetic speech on a syllable-level with the neural data time-series to obtain a target speech waveform for training the decoder. **e.** A representative example of causally synthesized speech from neural data, which matches the target speech with high fidelity.

We developed a real-time neural decoding pipeline (**Fig. 1c**) to synthesize T15’s voice instantaneously from intracortical neural activity, with continuous audio feedback, as he attempted to speak sentences cued on a screen at his own pace. Since the participant could not speak intelligibly, we did not have the ground truth for how and when he attempted to speak. Therefore, to generate aligned neural and voice data for training the decoder, we developed an algorithm to identify putative syllable boundaries directly from neural activity. This allowed us to generate target speech that was time-aligned to neural recordings as a proxy to T15’s intended speech (**Fig. 1d**).

We trained a multilayered Transformer-based^19^ model to causally predict spectral and pitch features of the target speech every 10 ms using the preceding binned threshold crossings and spike-band power. The base Transformer model architecture was augmented to compensate for session-to-session neural signal nonstationarities^20^ and to lower the inference time for instantaneous voice synthesis. The entire neural processing, from signal acquisition to synthesis of speech samples, occurred within 10 ms, enabling nearly-instantaneous speech synthesis (**Extended Data Fig. 2** shows end-to-end timings). The resulting audio was synthesized into voice samples by a vocoder^21^ and continuously played back to T15 through a speaker (**Fig. 1e**).

### Flexible and accurate closed-loop voice synthesis

We first tested the brain-to-voice BCI’s ability to causally synthesize voice from neural activity while T15 attempted to speak cued sentences (**Fig. 2a** and **Video 2**). Each trial consisted of a unique sentence which was never repeated in the training or evaluation trials. The synthesized voice was similar to the target speech (**Fig. 2b**), with a Pearson correlation coefficient of 0.89±0.04 across 40 Mel-frequency bands (**Extended Data Fig. 3a** reports Mel-cepstral distortion). We quantified intelligibility by asking 15 human listeners to match each of the 956 evaluation sentences with the correct transcript (choosing from 6 possible sentences of the same length). The mean and median accuracies were 94.34% and 100%, respectively (**Fig. 2i**). The instantaneous voice synthesis accurately tracked T15’s pace of attempted speech (**Extended Data Fig. 4**), which – due to his ALS – meant slowly speaking one word at a time. These results demonstrate that the real-time synthesized speech recapitulates the intended speech to a high degree, and can be identified by non-expert listeners. We also demonstrated that this brain-to-voice speech neuroprosthesis could be paired with our previously-reported high accuracy brain-to-text decoder^1^, which essentially acted as closed-captioning (**Video 3**).

**Fig. 2.**
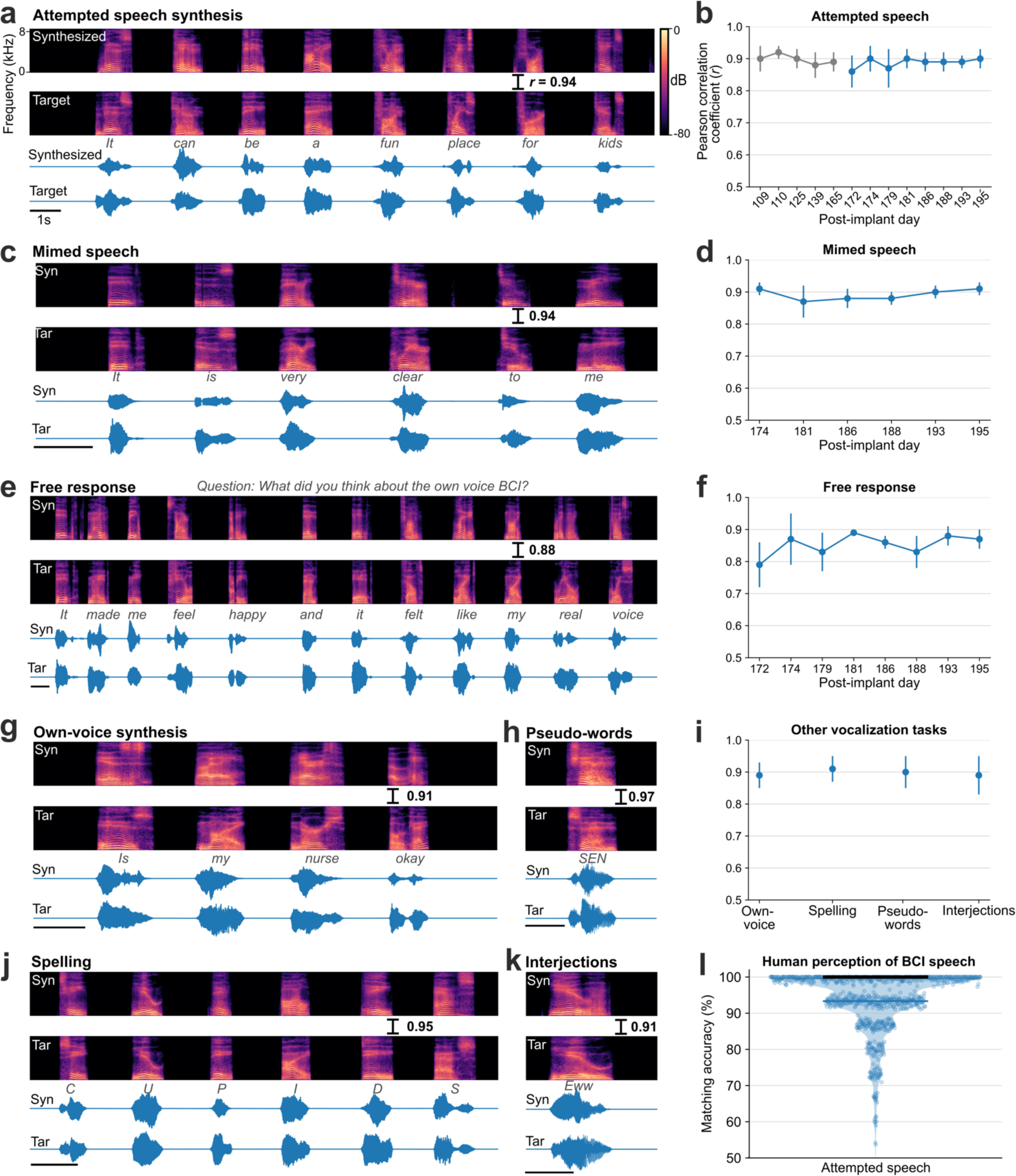
Voice neuroprosthesis allows a wide range of vocalizations. **a.** Spectrogram and waveform of an example trial showing closed-loop synthesis during attempted speech of a cued sentence (top) and the target speech (bottom). The Pearson correlation coefficient (*r*) is computed across 40 Mel-frequency bands between the synthesized and target speech. **b**. Pearson correlation coefficients (mean ± s.d) for attempted speech of cued sentences across research sessions. Sessions in blue were predetermined as evaluation sessions and all performance summaries are reported over these sessions. **c**. An example mimed speech trial where the participant attempted to speak without vocalizing and **d.** mimed speech Pearson correlations across sessions. **e.** An example trial of self-guided attempted speech in response to an open-ended question and **f.** self-guided speech Pearson correlations across sessions. **g**. An example personalized own-voice synthesis trial. **h, j, k.** Example trials where the participant said pseudo-words, spelled out words letter by letter, and said interjections, respectively. The decoder was not trained on these words or tasks. **i**. Pearson correlation coefficients of own-voice synthesis, spelling, pseudo-words and interjections synthesis. **l**. Human perception accuracy of synthesized speech where 15 naive listeners for each of the 956 evaluation sentences selected the correct transcript from 6 possible sentences of the same length. Individual points on the violin plot show the average matching accuracy of each evaluation sentence (random vertical jitter is added for visual clarity). The bold black line shows median accuracy (which was 100%) and the thin blue line shows the (bottom) 25^th^ percentile.

All four arrays showed significant speech-related modulation and contributed to voice synthesis, with the most speech-related modulation on the v6v and 55b arrays and much less speech-related modulation on the d6v array (**Extended Data Fig. 4**). Thanks to this high neural information content, the brain-to-voice decoder could be trained even with limited data, as shown by an online demonstration using a limited 50-word vocabulary on the first day of neuroprosthesis use (**Video 14**). Lastly, we compared this instantaneous voice synthesis method to an acausal method^3^ that decoded a sequence of discrete speech units at the end of each sentence (**Audio 1**). As expected, acausal synthesis – which benefits from integrating over the entire utterance – generated high quality voice (MCD 2.4±0.03); this result illustrates that instantaneous voice synthesis is a substantially more challenging problem.

People with neurodegenerative diseases may eventually lose their ability to vocalize all together, or may find vocalizing tiring. We therefore tested the brain-to-voice BCI during silent “mimed” speech where the participant was instructed to attempt to mouth the sentence without vocalizing. Although the decoder was only trained on attempted vocalized speech, it generalized well to mimed speech: the Pearson correlation coefficient was 0.89±0.03, which was not statistically different from voice synthesis during vocalized attempted speech (**Fig. 2c, d** and **Video 4**). **Extended Data Fig. 3b** shows human perception accuracy of synthesized speech during miming. T15 reported that he found attempting mimed speech less tiring compared to vocalized speech.

The aforementioned demonstrations involved T15 attempting to speak cued sentences. Next, we tested if the brain-to-voice BCI could synthesize unprompted self-initiated speech, more akin to how a neuroprosthesis would be used for real-world conversation. We presented T15 with questions on the screen (including asking for his feedback about the voice synthesis), which he responded to using the brain-to-voice BCI (**Fig. 2e**). We also asked him to say whatever he wanted (**Video 5**). The accuracy of his free response synthesis was slightly lower than that of cued speech (Pearson correlation coefficient 0.84±0.1, **Fig. 2f**, Wilcoxon rank-sum, *p*=10^-6^, n_1_=57, n_2_=933). We speculate that this reflected him using a different attempted speech strategy (with less attention to enunciating each phoneme) that he commonly used for his personal use with the brain-to-text BCI^1^.

This brain-to-voice decoder directly predicts acoustic speech features, which allows the user to produce a variety of expressive sounds, including non-word sounds and interjections, which are not possible with language- and vocabulary-dependent speech BCIs. To demonstrate this flexibility, we instructed T15 to use the brain-to-voice BCI to say made-up pseudo-words and interjections (e.g., “aah”, “eww”, “ooh”, “hmm”, “shoo”) (**Fig. 2h, k** and **Videos 7, 8**). The neuroprosthesis also enabled T15 to spell out words one letter at a time (**Fig. 2j** and **Video 9**). The brain-to-voice decoder was not trained on pseudo-words, spellings or interjections tasks but was able synthesize these sounds with a Pearson correlation coefficient of 0.90±0.01 (**Fig. 2i**).

Voice is an important element of people’s identities, and synthesizing a user’s own voice could further improve the restorative aspect of a speech neuroprosthesis. We therefore demonstrated that the instantaneous brain-to-voice framework was personalizable and could approximate T15’s pre-ALS voice (**Fig. 2g** and **Video 6**). To achieve this, we trained the brain-to-voice decoder on target speech produced by a voice cloning text-to-speech algorithm^22^ that sounded like T15. The participant used the speech synthesis BCI to report that listening to his own voice “*made me feel happy and it felt like my real voice*” (**Fig. 2e**). The accuracy of the own-voice synthesis was similar to the default voice synthesis (Pearson correlation coefficient of 0.87±0.04, **Fig. 2i**).

Through these varied speech tasks, we demonstrated that the brain-to-voice BCI framework is flexible and generalizable, enabling the participant to synthesize a wide variety of vocalizations.

### Online decoding of paralinguistic features from neural activity

Paralinguistic features such as pitch, cadence, and loudness play an important role in human speech, allowing us to be more expressive. Changing the stress on different words can change the semantic meaning of a sentence; modulating intonations can convey a question, surprise or other emotions; and modulating pitch allows us to sing. Incorporating these paralinguistic features into BCI-synthesized voice is an important step towards restoring naturalistic speech. We investigated whether these paralinguistic features are encoded in the neural activity in the ventral precentral gyrus and developed algorithms to decode and modulate these speech features during closed-loop voice synthesis.

Since the brain-to-voice decoder causally and immediately synthesizes voice, it inherently captures the natural pace of T15’s speech. To quantify this, T15 was asked to speak sentences at either a faster or slower speed. The voice synthesized by the neuroprosthesis reflected his intended speaking speed (**Fig. 3a**). **Fig. 3b** shows the differing distributions of durations of synthesized words attempted at fast (average speed of 0.97±0.19 s per word) and slow (average speed of 1.46±0.31 s per word) speeds. Additionally, we were able to decode quiet or loud attempted speech loudness from the neural features with 90% accuracy (**Extended Data Fig. 5**).

**Fig. 3.**
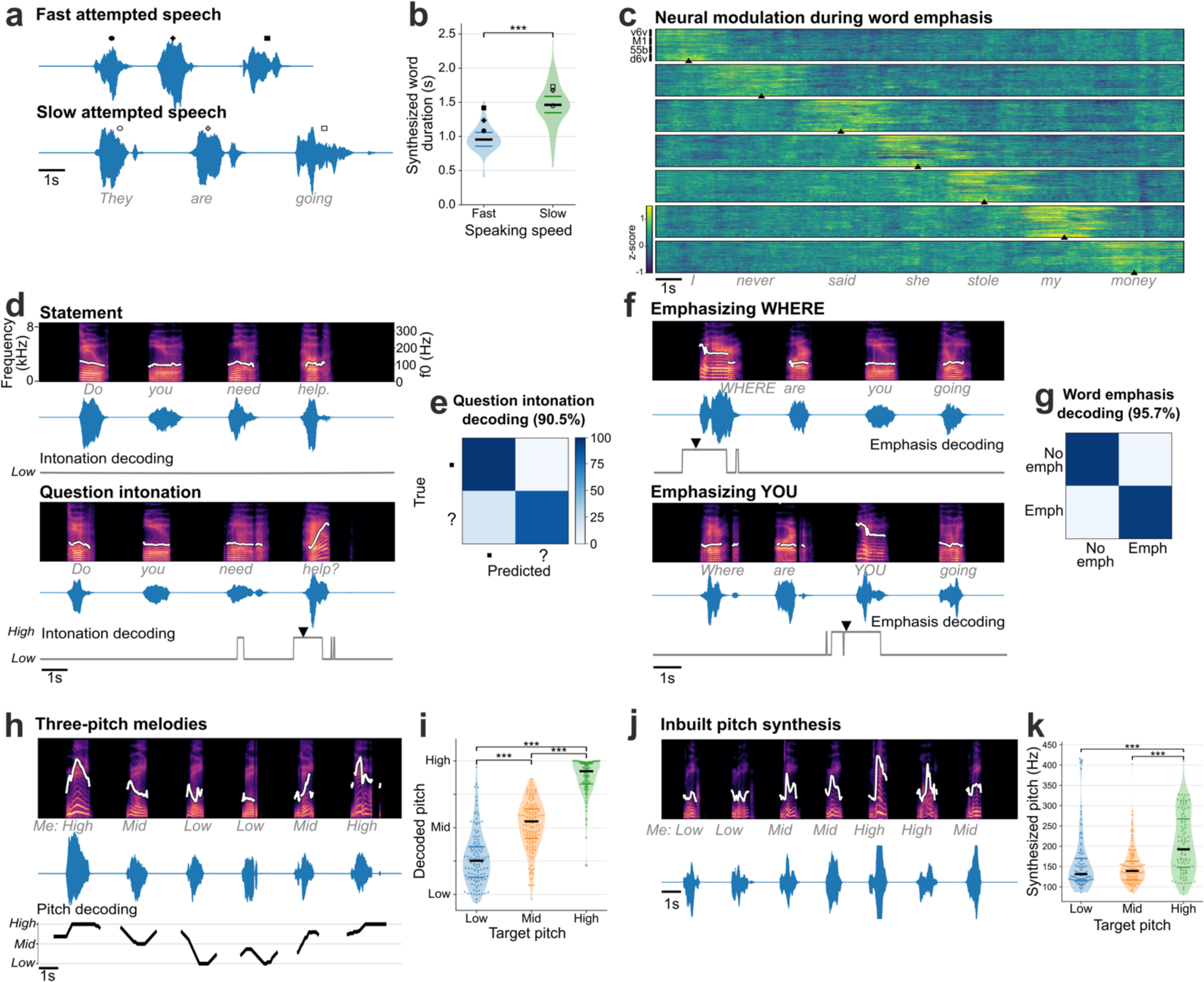
Modulating paralinguistic features in synthesized voice. **a.** Two example synthesized trials are shown where the same sentence was spoken at faster and slower speeds. **b**. Violin plots showing significantly different durations of words instructed to be spoken fast and slowly (Wilcoxon rank-sum, *p*=10^-14^, n_1_=72, n_2_=57). The bold black horizontal line shows the median value of the synthesized word duration and thin colored horizontal lines show the range between 25^th^ and 75^th^ percentiles. **c**. Trial-averaged normalized spike-band power (each row in a panel is one electrode) during trials where the participant emphasized each word in the sentence “*I never said she stole my money*”, grouped by the emphasized word. Trials were aligned using dynamic time warping and the mean activity across all trials was subtracted to better show the increased neural activity around the emphasized word. The emphasized word’s onset is indicated by the arrowhead at the bottom of each condition. **d**. Spectrograms and waveforms of two synthesized voice trials where the participant says the same sentence as a statement and as a question. The intonation decoder output is shown below each trial. An arrowhead marks the onset of causal pitch modulation in the synthesized voice. The white trace overlaid on the spectrograms shows the synthesized pitch contour, which is constant for a statement and increases during the last word for a question. **e**. Confusion matrix showing accuracies for closed-loop intonation modulation during real-time voice synthesis. **f**. Spectrograms and waveforms of two synthesized voice trials where different words of the same sentence are emphasized, with pitch contours overlaid. Emphasis decoder output is shown below. Arrowheads show onset of emphasis modulation. **g**. Confusion matrix showing accuracies for closed-loop word emphasis during real-time voice synthesis. **h**. Example trial of singing a melody with three pitch targets. The pitch decoder output that was used to modulate pitch during closed-loop voice synthesis is shown below. The pitch contour of the synthesized voice shows different pitch levels synthesized accurately for the target cued melody. **i.** Violin plots showing significantly different decoded pitch levels for low, medium and high pitch target words (Wilcoxon rank-sum, *p*=10^-14^ with correction for multiple comparisons, n_1_=122, n_2_=132, n_3_=122). Each point indicates a single trial. **j**. Example three-pitch melody singing synthesized by a unified brain-to-voice model. The pitch contour of the synthesized voice shows that the pitch levels tracked the target melody. **k**. Violin plot showing peak synthesized pitch frequency achieved by the inbuilt pitch synthesis model for low, medium and high pitch targets. Synthesized high pitch was significantly different from low and medium pitch (Wilcoxon rank-sum, *p*=10^-3^, n_1_=106, n_2_=113, n_3_=105). Each point shows an individual trial.

Next, we decoded the intent to modulate intonation to ask a question or to emphasize a specific word. We recorded neural activity while T15 attempted to speak the same set of sentences as either statements (no extra modulation in pitch) or as questions (with increasing pitch at the end of the sentence). This revealed increased neural activity recorded on all four arrays towards the end of the questions (**Extended Data Fig. 6**). To study the effect of attempted word emphasis on neural activity, in a different experiment we asked T15 to emphasize one of the seven words in the sentence “*I never said she stole my money*” by increasing that word’s pitch. This sentence, modeled after ^23^, changes its semantic meaning for each condition whilst keeping the phonemic content the same. Similar to the effect observed during the question intonation task, we observed increased neural activity around the emphasized word (**Fig. 3c**) on all four arrays (**Extended Data Fig. 7**) starting ∼350 ms prior to the onset of the word.

As a proof-of-principle that these paralinguistic features could be captured by a speech neuroprosthesis, we trained two separate binary decoders to identify the change in intonation during these question intonation and word emphasis tasks. We then applied these intonation decoders in parallel to the brain-to-voice decoder to modulate the pitch and amplitude of the synthesized voice in closed loop, enabling T15 to ask a question or emphasize a word (**Extended Data Fig. 8**). **Fig. 3d** shows two example closed-loop voice synthesis trials, including their pitch contours, where T15 spoke a sentence as a statement and as a question. The synthesized speech pitch increased at the end of the sentence during question intonation (**Video 10**). **Fig. 3f** shows two example synthesized trials of the same sentence where different words were emphasized (**Video 11**). Across all closed-loop evaluation trials, we decoded and modulated question intonation with 90.5% accuracy (**Fig. 3e**) and word emphasis with 95.7% accuracy (**Fig. 3g**).

After providing the aforementioned binary intonation control for questions or word emphasis, we investigated decoding multiple pitch levels from neural activity. We designed a three-pitch melody task where T15 attempted to “sing” different melodies consisting of 6 to 7 notes of low, medium and high pitch (e.g., low-mid-high-high-mid-low). These data were used to train a two-stage Transformer-based pitch decoder. During closed-loop voice synthesis, this pitch decoder ran simultaneously with the brain-to-voice decoder to modulate its pitch output; visual feedback of the decoded pitch level was also provided on-screen (**Video 12**). T15 was able to control the synthesized melody’s pitch levels (**Fig. 3h)**. **Fig. 3i** shows three distinct distributions of pitch levels decoded from neural activity across all singing task evaluation trials, demonstrating that the pitch and phonemic content of speech could be simultaneously decoded from neural activity in real-time.

In the preceding experiments, we used a data-efficient separate discrete decoder to modulate the synthesized voice because the vast majority of our training data consisted of neutral sentences without explicit instructions to modulate intonation or pitch. However, a more generalizable approach would be to develop a unified (single) brain-to-voice decoder that takes into account these paralinguistic features. We demonstrated the feasibility of such an approach by training our regular brain-to-voice decoder model architecture with the time-aligned target speech consisting of different pitch levels for target notes in the three-pitch singing task. This enabled the decoder to implicitly learn the mapping between neural features and the desired pitch level in addition to learning the mapping from neural activity to phonemic content (as before). During continuous closed-loop voice synthesis evaluation, this unified “pitch-enhanced” brain-to-voice decoder was able to synthesize different pitch levels as T15 attempted to sing different melodies (**Video 13** and **Fig. 3j, k**). This demonstrates that the brain-to-voice BCI framework has an inherent capability to synthesize paralinguistic features if provided with training data where the participant attempts the desired range of vocal properties (in this case, pitch).

### Rich output-null neural dynamics during speech production

Instantaneous brain-to-voice synthesis provides a unique view into neural dynamics with high temporal precision. We noticed that neural activity increased prior to and during the utterance of each word in a cued sentence, but that the aggregate neural activity decreased over the course of the sentence (**Fig. 1d, Extended Data Fig. 4**). Yet despite this broad activity decrease, the synthesis quality remained consistent throughout the sentence (**Extended Data Fig. 9)**. This seeming mismatch between overall neural activity and voice output suggested that the “extra” activity – which preceded voice onset for each word and also gradually diminished towards the end of a sentence – could be a form of output-null neural subspace activity previously implicated in movement preparation^24^, feedback processing^25^, and other computational support roles^26^.

We estimated the output-null and output-potent neural dimensions by linearly decomposing the population activity into a subspace that best predicted the speech features (output-potent dimensions, which putatively most directly relate to behavioral output) and its orthogonal complement (output-null dimensions, which putatively have less direct effect on the behavioral output). Both subspaces contained substantial speech-related information: the Pearson correlation coefficients of decoding speech using only the output-potent dimensions (which captured 2.5% of total variance) or only the output-null dimensions (97.5% of total variance) were 0.79±0.08 and 0.84±0.05, respectively. **Fig. 4a** shows output-null and output-potent components of neural activity around the onset of each word in sentences of different lengths. A clear decrease in output-null activity can be seen over the course of a sentence regardless of its length, whereas the output-potent activity remains consistent (**Fig. 4c**). An exception to this was the very last word, which tended to have an increase in output-null activity, especially as the last word was being finished. We do not know why the end of the sentence exhibited this effect but speculate that it is related to an end-of-trial cognitive change (e.g., the participant assessing his performance).

**Fig. 4.**
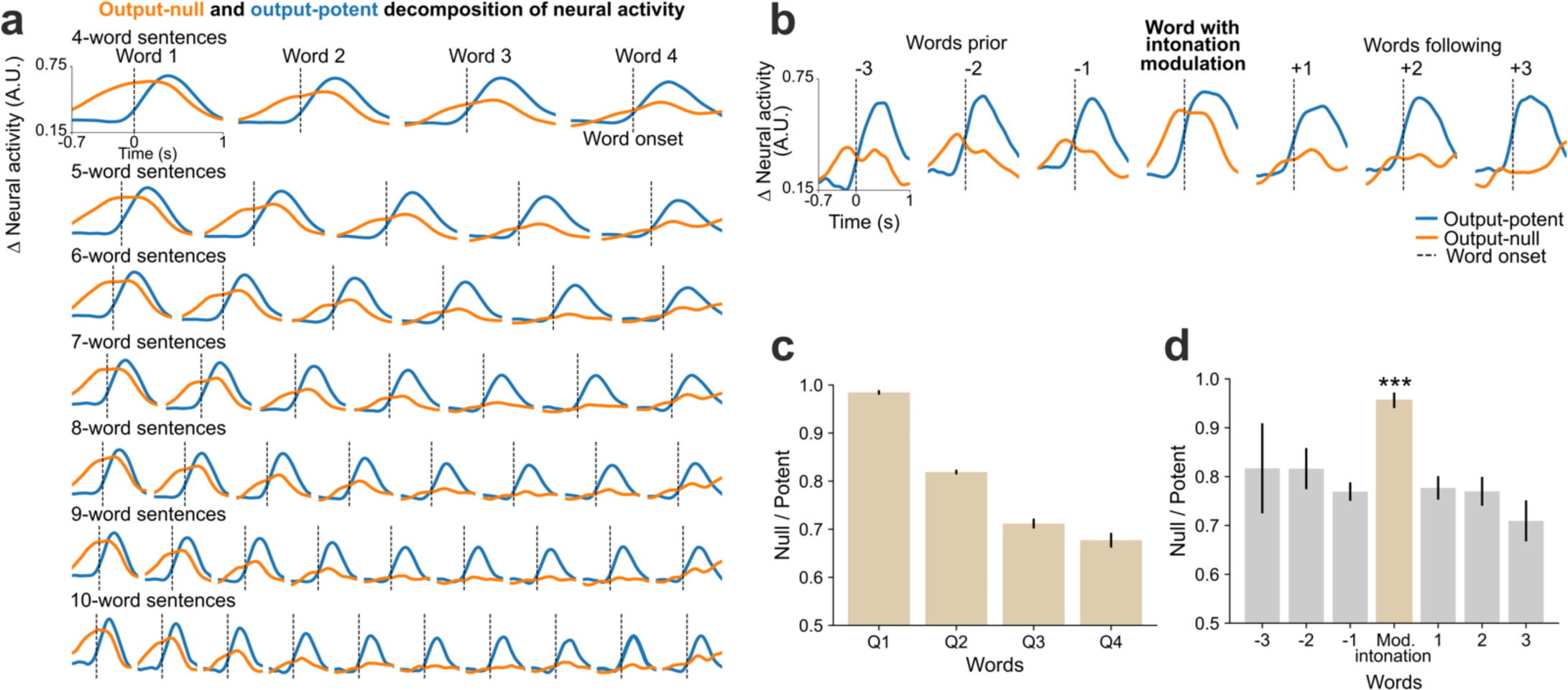
Output-null and output-potent neural dynamics during speech production. **a.** Average approximated output-null (orange) and output-potent (blue) components of neural activity during attempted speech of cued sentences of different lengths. Output-null activity gradually decayed over the course of the sentence, whereas the output-potent activity remained consistent irrespective of the length of the sentence. **b**. Average output-null and output-potent activity during intonation modulation (question-asking or word emphasis); data are trial-averaged aligned to the emphasized word (center) and the words preceding and/or following that word in the sentence. The output-null activity increased during pitch modulation as compared to the words preceding or following it. **c**. Panel **a** data are summarized by taking the average null/potent activity ratios for words in the first-quarter, second-quarter, third-quarter, and fourth-quarter of each sentence (mean ± s.e.). **d.** Panel **b** data are summarized by calculating average null/potent activity ratios of the intonation modulated word (beige) and the words preceding or following it (gray) (mean ± s.e.). The null/potent ratios of modulated words were significantly different from that of non-modulated words (Wilcoxon rank-sum, *p*= 10^-21^, n_1_=460, n_2_=922). **Extended Data Fig. 10** shows these analyses for each array individually.

We also examined the putative output-null and output-potent activity when the participant volitionally modulated his intonation. We found that the output-null activity increased significantly (*p*=10^-21^) for the word that was modulated (**Fig. 4b, d**) as compared to the words that preceded or followed it, explaining the previously-noted increase in overall neural activity preceding intonation-emphasized words (**Fig. 3c, Extended Data Fig. 6**).

## Discussion

This study demonstrated a “brain-to-voice” neuroprosthesis that directly mapped the neural activity recorded from four microelectrode arrays spanning ventral precentral gyrus into acoustic features. A man with severe dysarthria due to ALS used the system to synthesize his voice in real-time as he attempted to speak in both highly structured and open-ended conversation. The resulting voice was often intelligible. The decoding models were trained for a participant who could no longer speak intelligibly (and thus could not provide a ground truth speech target), and could be adjusted to emulate his pre-ALS voice. Unlike prior studies^3,17^, this brain-to-voice neuroprosthesis output sounds as soon as the participant tried to speak, without being restricted to a small number of words^17^ and without a constrained intermediate representation of discrete speech units that were generated after completion of each sentence^3^. To demonstrate the flexibility conferred by this direct voice synthesis BCI, the participant used it to synthesize various vocalizations including unseen words, interjections, and made-up words.

Furthermore, this study demonstrates that a brain-to-voice neuroprosthesis can restore additional communication capabilities over existing brain-to-text BCIs^1–3,27^. Neuronal activity in precentral gyrus encoded both phonemic and paralinguistic features simultaneously. Beyond providing a more immediate way to say words, this system could decode the neural correlates of loudness, pitch, and intonation. In online demonstrations, the neuroprosthesis enabled the participant to control a variety of aspects of his instantaneous digital vocalization, including the duration of words, emphasizing specific words in a sentence, ending a sentence as either a statement or a question, and singing three-pitch melodies. This represents a step towards restoring the ability of people living with speech paralysis to regain the full range of expression provided by the human voice.

We note that participant T15, who had been severely dysarthric for several years at time of this study, reported that he found it difficult to try to precisely modulate the tone, pitch, and amplitude of his attempted speech. Thus, we propose that using discrete classifiers to generate real-time modulated voice (which provides feedback to the participant that helps them mentally “hone in” on how to modulate their voice) can provide an intermediate set of training data useful for training a single unified decoder capable of continuous control of phonemic and paralinguistic vocal features. We demonstrated a proof of concept of this unified approach by training a single core decoder to intrinsically synthesize voice with different pitch levels, which the participant used for singing melodies.

A functional neuroanatomy result observed in this study that would not be predicted from prior ECoG^23,28–30^ and microstimulation studies^31,32^ is that the neural activity is correlated with paralinguistic features across all four microelectrode arrays, from ventral-most precentral gyrus to the middle precentral gyrus. We also observed that cortical activity across all four arrays increased well before attempted speech. We hypothesize that this reflects output-null preparatory activity^24,26^, and note that its presence is particularly fortuitous for the goal of causally decoding voice features because it gives the decoder a “sneak peek” shortly before intended vocalization. A particularly interesting observation was that this output-null activity seems to decrease over the course of a sentence. This may indicate that the speech motor cortex has a “buffer” for the whole sentence, which is gradually emptied out as the sentence approaches completion. We also observed an increase in output-null activity preceding words that were emphasized or modulated, which we speculate may be a signature of the additional neural computations involved in changing how that word is said. These results hint at considerable richness in speech-related motor cortical ensemble activity, beyond just the activity that is directly linked to driving the articulators. These phenomena represent an opportunity for future study, including leveraging the computation through dynamics framework and neural network modeling which have helped explain the complexity of motor cortical activity for preparing and producing arm and hand movements (reviewed in ^26^).

### Limitations

This study was limited to a single participant with ALS. It remains to be seen whether similar brain-to-voice performance will be replicated in additional participants, including those with other etiologies of speech loss. The participant’s ALS should also be considered when interpreting the study’s scientific results. Encouragingly, however, prior studies have found that neural coding observations related to hand movements have generalized across people with ALS and able-bodied animal models^33^ and across a variety of etiologies of BCI clinical trial participants^34,35^. Furthermore, the phonemic and paralinguistic tuning reported here at action potential resolution has parallels in meso-scale ECoG measurements over sensorimotor cortex in able speakers being treated for epilepsy^23,28^.

Although the performance demonstrated compares favorably with prior studies, the synthesized words were still not consistently intelligible. We also anecdotally observed that the participant’s energy level and engagement on a given block, as well as whether he attempted to enunciate the words clearly and fully, influenced synthesis quality. Brain-to-voice evaluations performed during the research sessions provided limited opportunity for practice-based improvement (i.e., sensorimotor learning). It remains an open question whether consistent long-term use will result in improved accuracy due to additional training data and/or learning. Separately, we predict that accuracy improvement is possible with further algorithm refinement and increasing the number of electrodes, which was previously shown to improve brain-to-text decoding accuracy^1,2^.

## Acknowledgements

We thank participant T15 and his family and care partners for their immense contributions to this research.

## Funding

Funded by: the Office of the Assistant Secretary of Defense for Health Affairs through the Amyotrophic Lateral Sclerosis Research Program under award number AL220043; a New Innovator Award (DP2) from the NIH Office of the Director and managed by NIDCD (1DP2DC021055); a Seed Grant from the ALS Association (23-SGP-652); A. P. Giannini Postdoctoral Fellowship (Card); Searle Scholars Program; a Pilot Award from the Simons Collaboration for the Global Brain (AN-NC-GB-Pilot Extension-00002343-01); NIH-NIDCD (U01DC017844); VA RR&D (A2295-R); Stavisky holds a Career Award at the Scientific Interface from BWF.

## Author Contributions

M.W., S.D.S., and D.M.B. conceived the study and experiment design. M.W. led the experiments, developed and implemented the target speech generation, decoder training algorithms, end-to-end pipeline for instantaneous speech synthesis: feature extraction, noise-removal, preprocessing, real-time brain-to-voice decoders, pitch decoders, vocoder and output audio playback, experimental tasks, post-processing, analyzed all the data and created figures. M.W. and N.S.C. developed and implemented the real-time neural signal processing, noise removal, and feature extraction pipelines. M.W., N.S.C., T.S-C., and X.H. coded the real-time data collection system and built the neuroprosthetic cart system. N.S.C. generated cloned voice samples for T15. M.W., N.S.C., and C.I. collected the primary data for this study. N.S.C. and C.I. interfaced with the participant and scheduled research sessions. D.M.B led planning and performing the surgical implant placement procedure. L.R.H. is the sponsor-investigator of the multisite clinical trial. D.M.B. was responsible for all clinical trial-related activities at UC Davis. S.D.S. and D.M.B. supervised all aspects of the project. M.W and S.D.S. wrote the manuscript. All authors reviewed and helped edit the manuscript.

## Competing Interests

Stavisky is an inventor on intellectual property owned by Stanford University that has been licensed to Blackrock Neurotech and Neuralink Corp. Wairagkar, Stavisky, and Brandman have patent applications related to speech BCI owned by the Regents of the University of California. Brandman is a surgical consultant to Paradromics Inc. Stavisky is a scientific advisor to Sonera and ALVI Labs. The MGH Translational Research Center has a clinical research support agreement with Neuralink, Synchron, Axoft, Precision Neuro, and Reach Neuro, for which Hochberg provides consultative input. Mass General Brigham (MGB) is convening the Implantable Brain-Computer Interface Collaborative Community (iBCI-CC); charitable gift agreements to MGB, including those received to date from Paradromics, Synchron, Precision Neuro, Neuralink, and Blackrock Neurotech, support the iBCI-CC, for which Hochberg provides effort.

## Data and materials availability

Data and code that can recreate the analyses and figures will be released with the peer-reviewed published paper.

## Methods

### Participant

A participant with amyotrophic lateral sclerosis (ALS) and severe dysarthria (referred to as ‘T15’), who gave informed consent, was enrolled in the BrainGate2 clinical trial (ClinicalTrials.gov Identifier: NCT00912041). This pilot clinical trial was approved under an Investigational Device Exemption (IDE) by the US Food and Drug Administration (Investigational Device Exemption #G090003). Permission was also granted by the Institutional Review Boards at the University of California, Davis (protocol #1843264) and Mass General Brigham (#2009P000505). T15 consented to publication of photographs and videos containing his likeness. This manuscript does not report any clinical trial-related outcomes; instead, it describes scientific and engineering discoveries that were made using data collecting in the context of the ongoing clinical trial.

T15 was a left-handed 45-year-old man. His ALS symptoms began five years before enrolment into this study. At the time of enrolment, he was non-ambulatory, had no functional use of his upper and lower extremities, and was dependent on others for activities of daily living (e.g., moving his wheelchair, dressing, eating, hygiene).

T15 had mixed upper and lower-motor neuron dysarthria and an ALS Functional Rating Scale Revised (ALSFRS-R) score of 23 (range 0 to 48 with higher scores indicating better function). He retained some neck and eye movements but had limited orofacial movement. T15 could vocalize but was unable to produce intelligible speech (see **Video 1**). He could be interpreted by expert listeners in his care team, which was his primary mode of communication.

Four chronic 64-electrode, 1.5 mm-length silicon microelectrode arrays coated with sputtered iridium oxide (Utah array, Blackrock Microsystems, Salt Lake City, Utah) were surgically placed in T15’s left precentral gyrus (putatively in the ventral premotor cortex, dorsal premotor cortex, primary motor cortex and middle precentral gyrus; **Fig. 1b**). The array placement locations were estimated based on pre-operative scans using the Human Connectome Project pipeline^1,18^. Voltage measurements from the arrays were transmitted to a percutaneous connection pedestal. An external receiver (Neuroplex-E, Blackrock Neuro) connected to the pedestal digitized and processed the measurements and sent information to a series of computers used for neural decoding. Data reported here are from post-implant days 25-342.

### Real-time neural feature extraction and signal-processing

Raw neural signals (voltage time series filtered between 0.3 to 7.5 kHz and sampled at 30 kHz with 250nV resolution) were recorded from 256 electrodes and sent to the processing computers in 1 ms packets. We developed the real-time signal processing and neural decoding pipeline using the custom-made BRAND platform^36^, where each processing step was conducted in a separate “node” running asynchronously.

We extracted neural features of action potential threshold crossings and spike-band power from each 1 ms incoming signal packet (30 samples) within 1 ms to minimize downstream delays. First, each packet was band pass filtered between 250 to 5000 Hz (4^th^ order zero-phase non-causal Butterworth filter) by adding 1 ms padding on both sides (from previous samples on one side and a constant mean value of the current samples on the other side) to minimize discontinuities at edges and denoised using Linear Regression Referencing (LRR)^37^. Then, threshold crossings were detected when the voltage dropped below -4.5 times the root mean squared (RMS) value for each channel (electrode). Spike-band power was computed by squaring and taking the mean of the samples in the filtered window for each channel and was clipped at 50k μV^2^ to avoid outliers. Neural features were binned into 10 ms non-overlapping bins (counting threshold crossings and taking the mean spike-band power across 10 consecutive feature windows, such that each of the 256 electrodes contributed two features). Each bin was first log-transformed, then normalized using rolling means and s.d. from the past 10 s for that feature. Each feature was then causally smoothed using a sigmoid kernel of length 1.5 s of the past activity. Thus, a vector of 1×512 binned neural features was sent to the brain-to-voice decoder every 10 ms. After each “block” of neural recording (a contiguous period of task performance), we re-computed RMS thresholds and LLR weights to be used in the next block. This helped in minimizing nonstationarities in the neural signals.

### Experimental paradigm

This study comprises multiple closed-loop speech tasks used to develop and evaluate a voice synthesis neuroprosthesis. Research sessions were structured as a series of blocks of ∼50 trials of a specific task. Each trial began with a “delay” period of 1.5-4 s in which a red square and a text cue was shown on the screen During this period, the participant was instructed to read the cue and prepare to speak. This was followed by a “go” period (indicated by a green square) where the participant was instructed to attempt to speak the cued text at his own pace after which he ended the trial using an eye tracker by looking at a “Done” icon on the screen. Closed-loop instantaneous voice synthesis was active during the “go” period. There was then a short 1-1.5 s interval before the start of the next trial.

The participant performed the following speech tasks using the above trial structure: 1) attempting to speak cued sentences; 2) miming (without vocalizing) cued sentences; 3) responding to open-ended questions in his own words or saying anything he wanted; 4) spelling out words letter by letter; 5) attempting to speak made-up pseudo-words; 6) saying interjections; 7) speaking cued sentences in fast or slow speeds; 8) speaking quietly or at a normal loudness; 9) modulating intonation to say a sentence as a statement or as a question; 10) emphasizing certain words in a sentence; and 11) singing melodies with different pitch level targets (this task had a prompted reference audio cue for the melody which was played during the delay period).

After the initial eight research sessions with a mix of open-loop and closed-loop blocks, all cued sentence trials in the rest of the sessions were conducted with either closed-loop voice synthesis, closed-loop text decoding^1^ or both to improve participant’s engagement in the task. All other types of tasks had closed-loop voice synthesis feedback. In a typical research session, we recorded ∼150-350 structured trials.

### Closed-loop continuous brain-to-voice synthesis

#### Target speech generation for decoder training

Since T15 was unable to produce intelligible speech, we did not have a ground-truth reference of his speech to match with the neural activity required for training a decoder to causally synthesize voice. Hence, we generated a “target” speech waveform aligned with neural activity as an approximation of T15’s intended speech.

We first generated synthetic speech waveforms from the known text cues in the training data using text-to-speech algorithm (native TTS application on MacOS version 13.5.1). Next, we identified putative syllable boundaries of T15’s attempted speech from the corresponding neural activity and aligned the synthetic speech by dynamically time stretching it to match these syllable boundaries to obtain the time-aligned target speech. The target speech was aligned at syllable level because syllables are the fundamental units of prosody in human speech^38^. During our first research session, there was no prior neural data available, so we used coregistered microphone recordings of T15’s attempted (unintelligible) speech to segment word boundaries and generate time-aligned target speech (i.e., unlike for later sessions, the session 1 time-alignment was done at a word rather than syllable level). This was done programmatically by identifying word boundaries based on the voice amplitude envelope of the participant’s attempted speech waveform, which was then adjusted manually if required. In subsequent sessions, we relied solely on neural data to estimate syllable boundaries: we used a brain-to-voice model trained on past neural data to synthesize speech and used its envelope to automatically segment syllable boundaries^39^, which were manually reviewed and occasionally adjusted if required. We then used dynamic time warping to align the envelope of the synthetic TTS speech to the envelope of the neurally predicted speech to create a mapping between their syllable boundaries. Then, the TTS segment within each syllable boundary was time-stretched to have the same duration as the corresponding segment of neurally predicted speech. This produced the target speech, which was now time-aligned with neural activity and suitable for training the new brain-to-voice decoder for the current session. This process was repeated iteratively for each session.

#### Brain-to-voice decoder architecture

The core brain-to-voice model was adapted from the Transformer architecture^19^. The model had two main components: an input embedding network and a base Transformer. Separate input embedding networks consisting of 2 fully-connected dense layers (512 and 128 units respectively, ReLU activation) were used for each week of neural data recording to compensate for week-to-week nonstationarities. The output from the input embedding network was passed into the base Transformer model consisting of 8 Transformer encoder blocks (head size 128, number of heads 4, a dropout of 0.5 after multi-head attention layer, each of the two feed-forward layers with 256 and 128 units respectively, and a normalization layer at the beginning and the end). Positional encoding was added before the first Transformer block. Additionally, we included residual connections between each Transformer block (separate from the residual connections within each block). The output sequence from Transformer blocks was pooled by averaging and passed to two dense layers (1024, 512 units, ReLU activation) and finally through a dense layer of size 20 to output 20-dimensional predicted speech features.

At each step, an input to the brain-to-voice decoder was a 600 ms window of binned neural features (threshold crossings and spike-band power) of shape 60×512 (60 bins of 10 ms with 256 channels x 2 features). The first layer of the model averaged two adjacent bins of the input sequence to reduce the sequence length by half whilst preserving the temporal information. The output of the decoder was a vector of 20 predicted speech features (which were then sent to a vocoder to generate synthesized speech samples in closed-loop blocks, described later). The decoder ran every 10 ms to produce a single 10 ms frame of voice samples. All the model hyperparameters were tuned manually with special consideration given to minimize the inference time for instantaneous closed-loop voice synthesis.

#### Decoder training

We trained a new decoder for each session using all cued sentence trials (which were unique) from all previous research sessions. To train the decoder robustly, we used 4-20 augmented copies of each trial. Neural features were augmented using three strategies: adding white noise (mean 0, s.d. 1.2) to all timepoints of all channels independently, a constant offset (mean 0, s.d. 0.6) to all spike-band channels independently and its scaled version (x0.67) to threshold crossings, and the same cumulative (random walk) noise (mean 0, s.d. 0.02) to all channels along the time course of the trial. We extracted a 600 ms sliding window (shifted by 10 ms from step to step) from continuous neural features and its corresponding 10 ms frame of output target speech features (20-dimensional vector) as a single training sample. These 20-dimensional output speech features (18 Bark cepstral coefficients, pitch period and pitch strength) for every 10 ms of target speech waveform were extracted from the time-aligned TTS target voice using the encoder for the pretrained LPCNet vocoder^21^. Each acoustic feature in the 20-dimensional vector was normalized independently based on the min and max of that feature (obtained from the training TTS dataset) to bring different acoustic features to the same scale (having all output features have a similar numeric range helps with accurate prediction). The features were then temporally smoothed before sending this vector as output in a batch for training the decoder.

The model was trained for ∼15-20 epochs with a batch size of 1024 samples (each epoch had ∼50k batches), a constant learning rate of 5×10^-4^, Adam optimizer (β1=0.9, β2=0.98, ε=1e-9) and Hubert loss (δ=1.35) which affords the advantage of both L1 and L2 losses and is less sensitive to outliers. The training took between 20 and 40 hrs on three NVIDIA GeForce RTX 3090 GPUs depending on the amount of data used for training.

On the first session of neural recording, we collected 190 open-loop trials of attempted speech from a 50-word vocabulary to train the decoder and were able to synthesize voice in closed-loop with audio feedback on the same day. Although the closed-loop synthesis on this data was less intelligible due to the model not being optimized on the first day, we later demonstrated offline that with an optimized model, we could achieve intelligible synthesis with this small amount of neural data and a limited vocabulary (**Video 14**).

In subsequent sessions, we collected more attempted speech trials with a large vocabulary and iteratively optimized our brain-to-voice decoder architecture to improve the synthesis quality. Here, we report the performance of closed-loop voice synthesis from neural activity using the “final” brain-to-voice decoder architecture for predetermined evaluation sessions. For each of these sessions, the decoder was trained on all the data collected up to one week prior (total of ∼5500-8900 trials).

For training the personalized-own voice synthesis model, we first generated time-aligned target speech that sounded like T15’s pre-ALS voice using the StyleTT2 text-to-speech model^22^ fine-tuned on T15’s voice samples prior to developing ALS. The rest of the process for decoder architecture and training was the same as above.

#### Closed-loop voice synthesis

During closed-loop real-time voice synthesis, we first extracted neural features every 1 ms. These were then binned, log-transformed, causally normalized and smoothed and aggregated into 600 ms causal sliding windows. This neural feature sequence was decoded by the brain-to-voice model into 20 acoustic speech features at each time step as described above. Inference was done on a single NVIDIA RTX A6000 GPU. The predicted speech features were rescaled back to their original range before normalization based on previously computed min and max of each feature (recall that during training, each acoustic speech feature was normalized independently such that all features were on the same scale and thus, the brain-to-voice decoder predicted normalized speech features which therefore needed rescaling for synthesis by the vocoder). The LPCNet vocoder requires 16 additional linear predictive coding (LPC) features in addition to the 20 acoustic features, but these are not independent and were derived from the 18 predicted cepstral features that were amongst the 20 decoded acoustic features. This resulted in a 36-feature vector (20 +16), which was synthesized into a single 10 ms frame of speech waveform (sampled at 16 kHz) using the pretrained LPCNet vocoder every 10 ms. The entire pipeline from neural signal acquisition to reconstruction of speech samples of a single frame took less than 10 ms (**Extended Data Fig. 2**). These samples were sent to the audio playback computer as they were generated, where they were played through a speaker continuously, thereby providing closed-loop audio feedback to the participant. We focused our engineering efforts on reducing the inference latency, which fundamentally bounds speech synthesis latency. However, pragmatically, we found that the largest latency occurred due to the audio playback driver (**Extended Data Fig. 2**). We were able to subsequently substantially lower these playback latencies and predict that further reductions are possible with additional optimization in interfacing with sound drivers. All results reported in this study are for closed-loop voice synthesis unless specified otherwise.

#### Evaluation of synthesized speech

We evaluated synthesized speech by measuring the Pearson correlation coefficient between the synthesized and TTS-derived target (fully intelligible) speech. We computed average Pearson correlations across 40 Mel-frequency bands of audio sampled at 16 kHz. The Mel-spectrogram with 40 Mel-frequency bands was computed using sliding (Hanning) windows of 50 ms with 10 ms overlap and converted to decibel units. We also computed Mel-cepstral distortion between the synthesized speech and the target speech using the method described in ^27^.

To evaluate human perception of BCI-synthesized speech, we asked 15 naïve listeners to listen to each synthesized speech trial and identify the transcript that matched the audio from six possible sentences of the same length. We used the crowd analytics platform Amazon Mechanical Turk to evaluate 1,014 synthesized sentences (vocalized and mimed trials) each by 15 individuals. To test if the crowd workers actually listened to the audio, we included a fully intelligible control audio clip with each synthesized audio. We rejected any trials with wrong answers for the control audio and resubmitted these trials for evaluation until we had 15 accepted answers per evaluation sentence.

### Decoding paralinguistic features for modulating synthesized voice

#### Decoding intonation intent from neural activity

We collected task blocks where T15 was instructed (1) to modulate his attempted speech intonation to say cued sentences as statements (no change in pitch) or (2) as questions (by changing the pitch from low to high towards the end of the sentence) or (3) to emphasize capitalized words in cued sentences (by increasing pitch with slight increase in loudness for emphasis). We analyzed question and word emphasis tasks separately but followed the same decoding procedure. We did not have the ground truth of when exactly T15 modulated his intonation to train the intonation decoders. Hence, trials were grouped by the cue sentence and their neural data aligned using dynamic time warping^40^. The average of these aligned trials was subtracted from each trial to reveal changes in neural activity (example, **Fig. 3c**). These trial-averaged data were used to manually label segments of neural data from each (warped) single trial as intonation modulation (class 1) or no modulation (class 0) for subsequent use in training intonation decoders.

For intonation decoding, we only used the spike-band power feature due to its higher signal-to-noise ratio (eliminating the threshold crossings helped reduce the feature size, which was helpful for this more data-limited decoder training). Sliding windows of 600 ms shifted by 10 ms were derived from binned neural activity to generate samples for decoder training. For each window, we took the mean of two adjacent bins to reduce the sequence length in half (to 30 bins from 60 bins), whilst preserving the temporal information. This sequence of features was then flattened to obtain a single 7,680-dimensional feature vector (30 x 256) as input to the two-class decoder. A binary logistic regression decoder was trained to classify the neural feature vectors into ‘no change in intonation’ (0) or ‘change in intonation’ (1). During closed-loop trials, this intonation decoder ran in parallel to the brain-to-voice decoder, and was used to predict intonation from features from the preceding 600 ms of neural activity, every 10 ms. Separate binary decoders were trained to detect intonation modulation for asking questions and for word emphasis. For a given task (questions or word emphasis), the intonation decoders ran simultaneously with the main brain-to-voice decoder.

#### Closed-loop intonation modulation in the synthesized voice

One of the speech features predicted by the brain-to-voice decoder characterizes the pitch component, which is used by the LPCNet vocoder to synthesize a speech waveform. To adjust the synthesized voice’s intonation in real-time, we artificially modified this feature’s value (relative to what the brain-to-voice decoder output) upon detection of a change in intonation from neural activity by the parallel intonation decoder. For the question intonation task, when the binary intonation decoder detected the question intonation (defined as when more than 60% of bins in the previous 700 ms window were classified as positive), it sent a trigger to modulate the pitch feature predicted by the brain-to-voice decoder according to a predefined pitch profile for asking a question (gradually increasing the pitch of the word from low to high). Each intonation modulation trigger was followed by a refractory period of 1.5 s to avoid consecutive duplicate triggers. The (now modified) speech features were then synthesized by the vocoder as previously described (see **Extended Data Fig. 8**).

Similarly, for emphasizing certain words in a sentence in closed-loop, the binary emphasis decoder sent a trigger (defined as when more than 60% of bins in the previous 800 ms window were classified as positive) to modulate pitch features predicted by the brain-to-voice decoder according to a predefined pitch profile for word emphasis – modulating the pitch from high to low – and increasing the volume of synthesized speech by 20%. We intentionally chose pitch profiles that were exaggerated so that the participant could clearly tell whether or not the intention modulation was working during the closed-loop tasks. However, these can trivially be adjusted in the future to avoid the resulting somewhat artificial-sounding exaggerated intonation.

We computed the accuracy of closed-loop intonation modulation by calculating what fraction of individual words in a sentence were modulated appropriately (i.e., matching the prompt).

#### Loudness decoding

The same architecture as for decoding intonation was followed to train a loudness decoder to classify neural data as belonging to quiet (class 0) speaking and normal speaking (class 1) from neural activity. Here the data labels were based on the task instruction. This decoder was evaluated offline by randomizing all the trials and splitting them into 13% heldout evaluation trials and 87% training trials.

#### Discrete pitch decoding for singing melodies

We collected neural data while T15 attempted to sing three-pitch melodies comprised of 6-7 notes of three pitch levels (e.g., low-mid-high-high-mid-low). Each note consisted of a short word to be sung in the target pitch level. At the start of each trial, an audio cue of the melody was played during the delay period for reference only. However, we did not instruct the participant to match the frequencies of the three notes used in these cued melodies because it was difficult for T15 to precisely modulate his pitch due to his severe dysarthria. Rather, he was instructed to try to make three distinct “low”, “medium”, and “high” pitch notes of his own choice.

We used a two-stage pitch decoding approach to decode pitch intent from spike-band power. A first Transformer-based decoder (same architecture as above, but with only two Transformer blocks and no input embedding network) was used to identify the participant’s intention to speak (i.e., classify between silence and intent to sing, based on training labels where ‘intent to sing’ was from 600 ms before the start of a note to the end of the note). A second Transformer-based decoder then decoded his intended pitch level (1-low, 2-mid, 3-high) if (and only if) an intention to sing was detected. Both decoders were trained using the categorical cross entropy loss. Since the results of the previous intonation modulation tasks showed that changes in paralinguistic features can be detected in advance of speech onset, labels for each pitch level were assigned to the neural data from 600 ms prior to the note onset to the end of the note attempted at that pitch.

During the closed-loop singing task, this two-staged pitch decoder ran simultaneously with the core brain-to-voice decoder. The output of the pitch decoder was smoothed with a moving average of a prior 800 ms window and then used to continuously modulate the predicted brain-to-voice pitch feature in real-time, which was then vocoded as usual (see **Fig. 3h, i** and **Extended Data Fig. 8**). Thus, the participant was able to sing melodies consisting of both phonemic content and three different pitches (e.g., “_la_ la ^la^”) through his synthesized voice. Additionally, we provided a closed-loop visual feedback on the screen by showing the decoded pitch level and interactive target cues for the note in the melody that T15 was singing (**Video 13**).

#### Inbuilt continuous pitch synthesis for singing melodies

In the previous intonation and pitch modulation tasks, we used a separate decoder to detect changes in the paralinguistic features and modulate the synthesized voice accordingly. Here we developed a *unified* brain-to-voice decoder that is inherently capable of synthesizing pitch in the melody singing task. To achieve this, we trained the regular brain-to-voice decoder now using neural data and target speech waveforms with varying pitch levels from the three-pitch singing task. These target waveforms were generated by first using the same time-aligned target speech generation algorithm described above from the TTS of the phonemic content of the note, and then adjusting the pitch to match the instructed pitch levels (low, mid, or high) of the notes in the cued melody (**Fig. 3j**).

### Output-null and output-potent analysis of neural activity

To study the underlying neural dynamics of speech production, we decomposed the neural activity into two orthogonal output-null and output-potent components using methods adapted from^25^. To do this, we first adopted a simplified linear decoding approach. We fit a linear decoder *y* = *Wx*, where *x* is a vector of neural features, *y* is a 20-dimensional vector of speech features, and *W* is the linear decoder, using ordinary least-squares regression. We trained a separate linear decoder for each session to account for session-to-session nonstationarities. The linear decoder matrix *W* was then decomposed into orthogonal null- and row-subspaces. The neural activity *x* was projected onto the null space (putative output-null dimensions of the neural activity) and row space (putative output-potent dimensions of the neural activity). The change in neural activity for null and row space projections for each trial was obtained by computing the Euclidean distance of the projections from the baseline activity (first 500 ms of the trial) and normalizing it between 0 and 1 (min and max across that whole trial) to get output-null and output-potent components, respectively. This normalization was used to account for potential neural nonstationarities across a session and between datasets, since ultimately we were interested in the relative changes of these neural projections over the course of each sentence.

Trial averaged output-null and output-potent neural activity components were obtained for all sentences of the same length between -700 ms to +1s from the onset of each word in a sentence (**Fig. 4**). This output-null and output-potent analysis was also performed on intonation modulation for questions and word emphasis tasks, and the output was compared with that of the regular cued attempted speech task.

### Statistical testing

We used a two-sided Wilcoxon rank-sum test to compare two groups of data. The *p*-values were corrected for multiple comparisons using Bonferroni correction where necessary. We used a non-parametric test because datasets being compared were of different size and normal distribution was not assumed because the actual underlying distribution was unknown.

## Extended data figures

**Extended Data Fig. 1:**
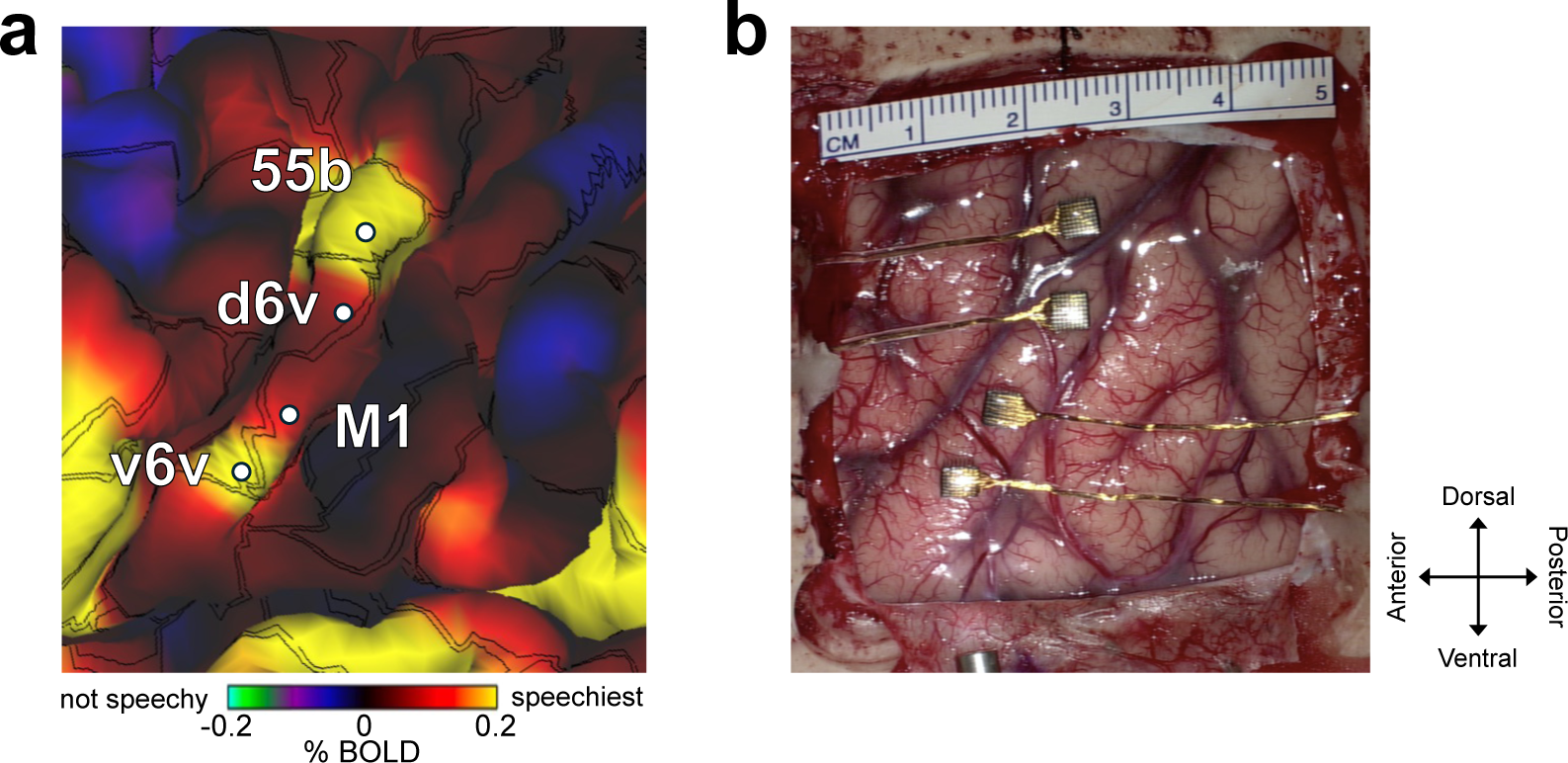
Microelectrode array placement. **a.** The estimated resting state language network from Human Connectome Project data overlaid on T15’s brain anatomy. **b**. Intraoperative photograph showing the four microelectrode arrays placed on the of T15’s precentral gyrus.

**Extended Data Fig. 2:**
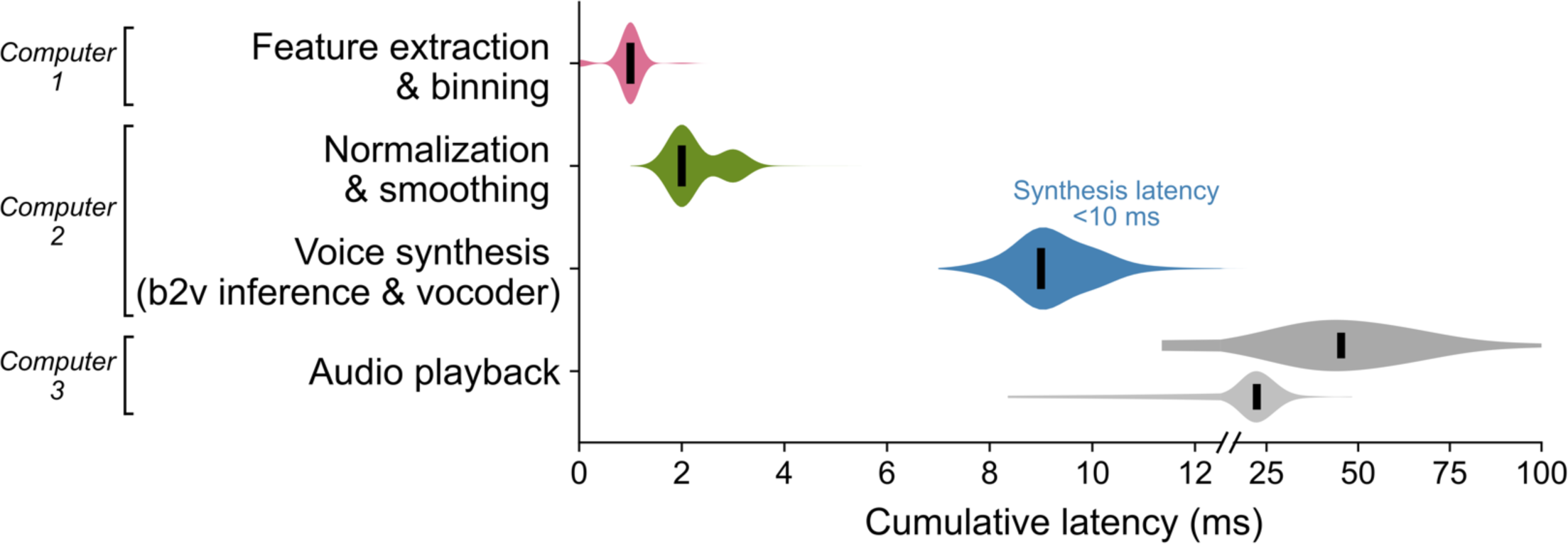
Latencies of closed-loop brain-to-voice synthesis. Cumulative latencies across different stages in the voice synthesis and audio playback pipeline are shown. Voice samples were synthesized from raw neural activity measurements within 10 ms and the resulting audio was played out loud continuously to provide closed-loop feedback. Note the linear horizontal axis is split to expand the visual dynamic range. We focused our engineering primarily on reducing the brain-to-voice inference latency, which fundamentally bounds the speech synthesis latency. As a result, the largest remaining contribution to the latency occurred after voice synthesis decoding during the (comparably more mundane) step of audio playback through a sound driver. The cumulative latencies with the audio driver settings used for T15 closed-loop synthesis are shown in dark gray. Audio playback latencies were subsequently substantially lowered through software optimizations (light gray) and we predict that further reductions will be possible with additional computer engineering.

**Extended Data Fig. 3:**
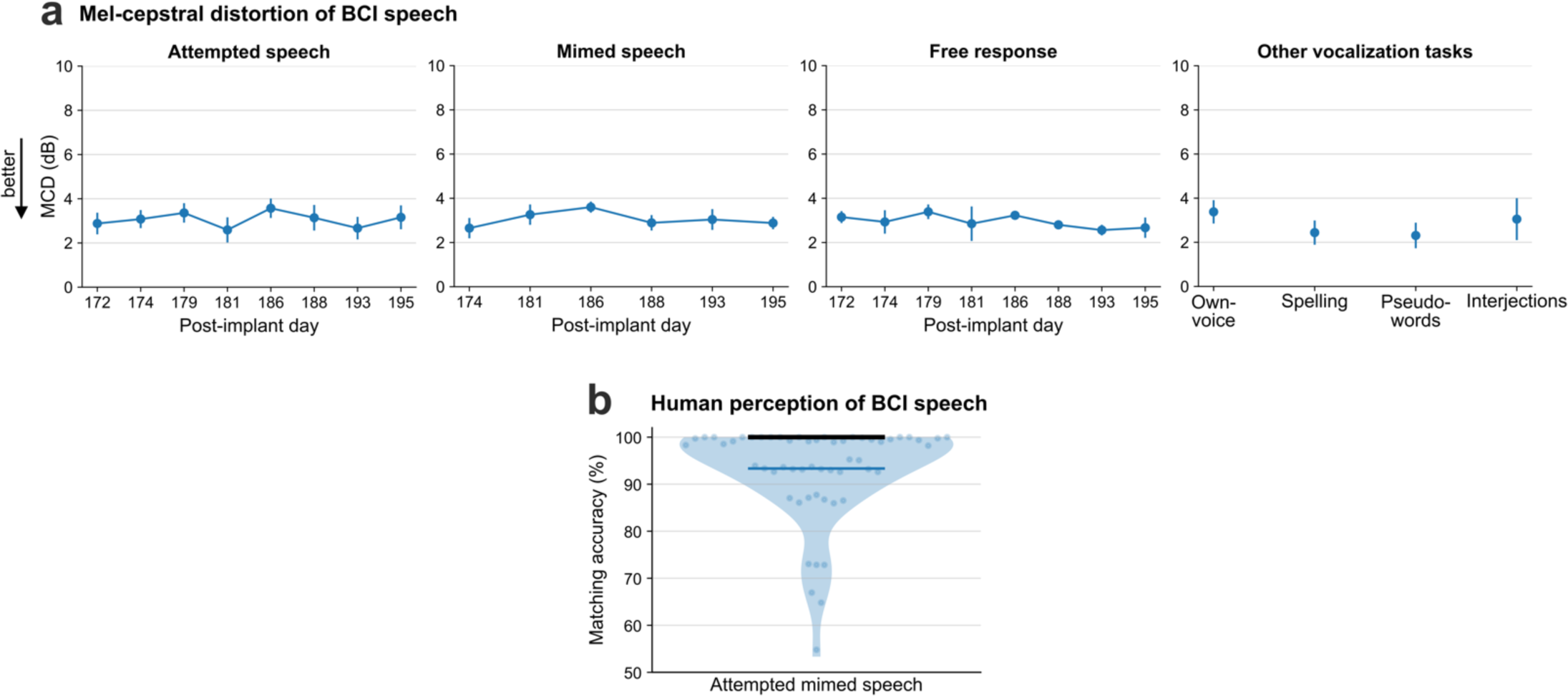
Additional BCI speech synthesis performance metrics. **a.** Mel-cepstral distortion (MCD) is computed across 25 Mel-frequency bands between the closed-loop synthesized speech and the target speech. The four subpanels show MCDs (mean ± s.d) between the synthesized and target speech for different speech tasks in evaluation research sessions. **b.** Human perception accuracy of BCI synthesized voice during mimed speech trials. 15 naïve listeners selected the transcript matching the synthesized speech from 6 possible sentences of the same length for each of the 58 evaluation sentences. Individual points on the violin plot show the average matching accuracy of each evaluation sentence (random small vertical jitter is added for visual clarity). The bold black line shows median accuracy (which was 100%) and the thin blue line shows the (bottom) 25^th^ percentile.

**Extended Data Fig. 4:**
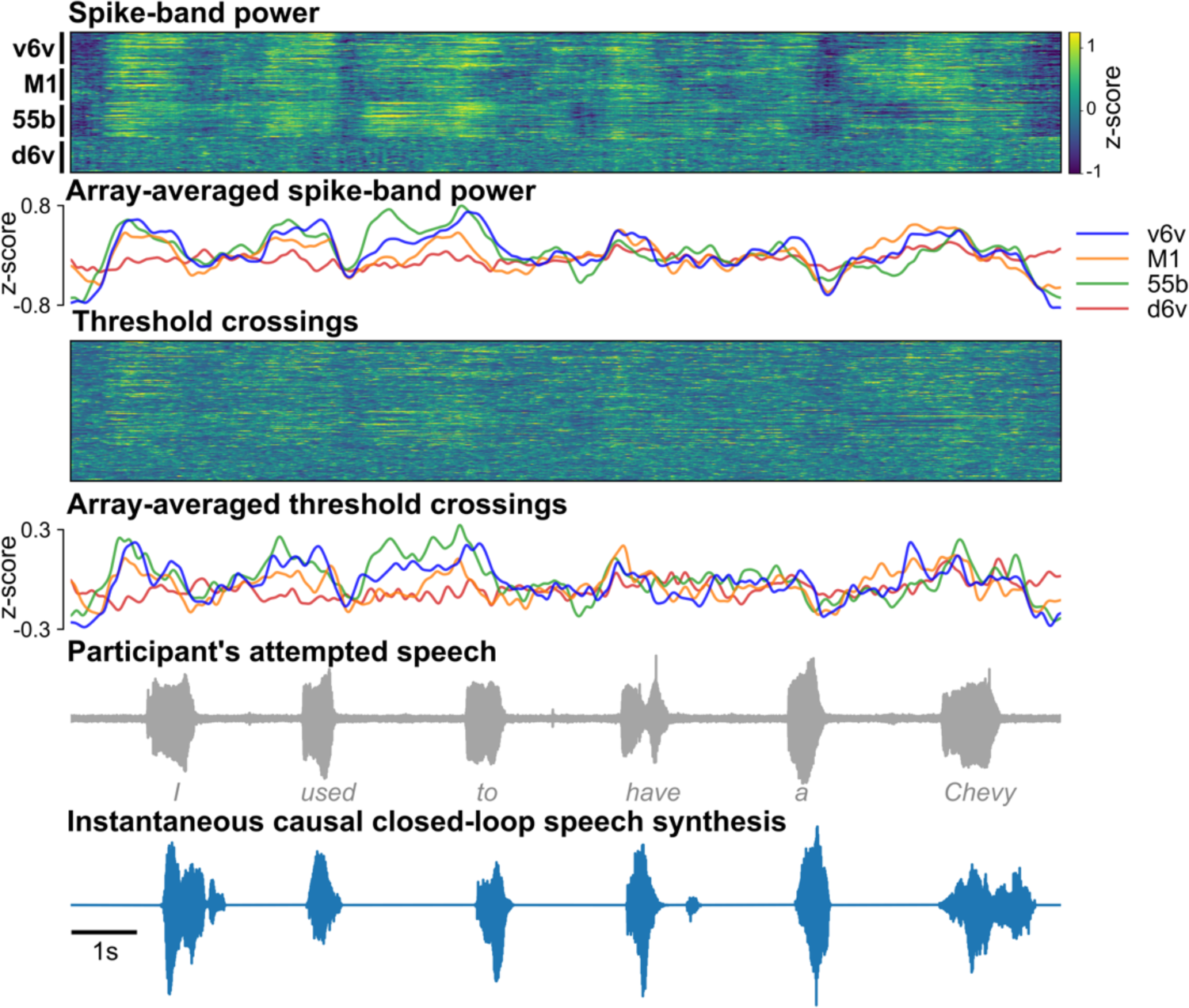
Example closed-loop speech synthesis trial. Spike-band power and threshold crossing spikes from each electrode are shown for one example sentence. These neural features are binned and causally normalized and smoothed on a rolling basis before being decoded to synthesize speech. The mean spike-band power and threshold crossing activity for each individual array are also shown. Speech-related modulation was observed on all arrays, with the highest modulation recorded in v6v and 55b. The synthesized speech is shown in the bottom-most row. The gray trace above it shows the participant’s attempted (unintelligible) speech as recorded with a microphone.

**Extended Data Fig. 5:**
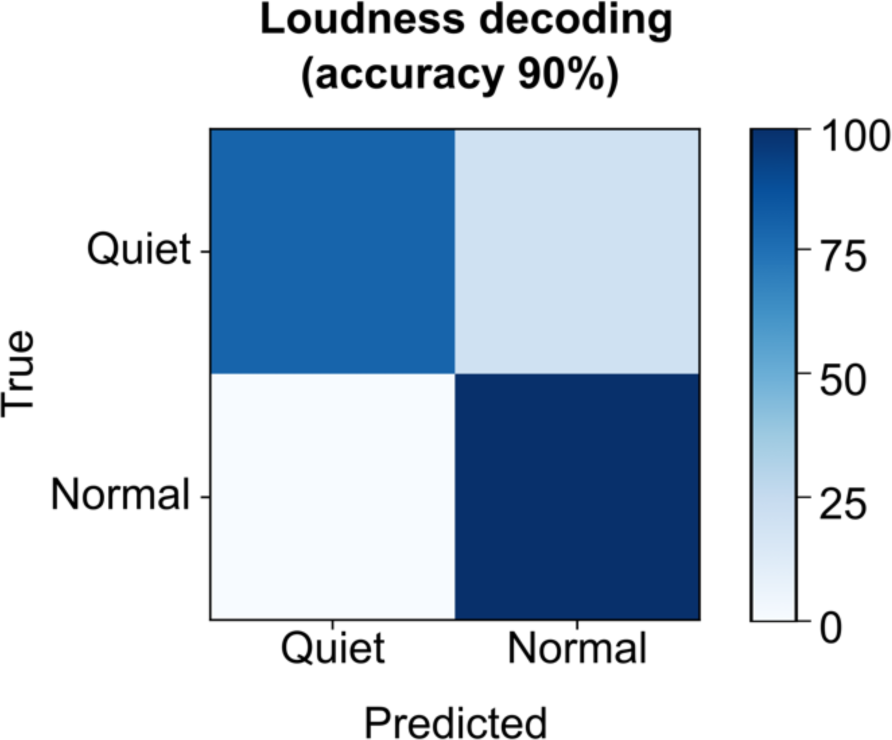
Loudness decoding from neural activity. Confusion matrix showing offline accuracies for classifying the loudness of attempted speech from neural activity using a binary decoder while the participant was instructed to speak either quietly or in his normal volume.

**Extended Data Fig. 6:**
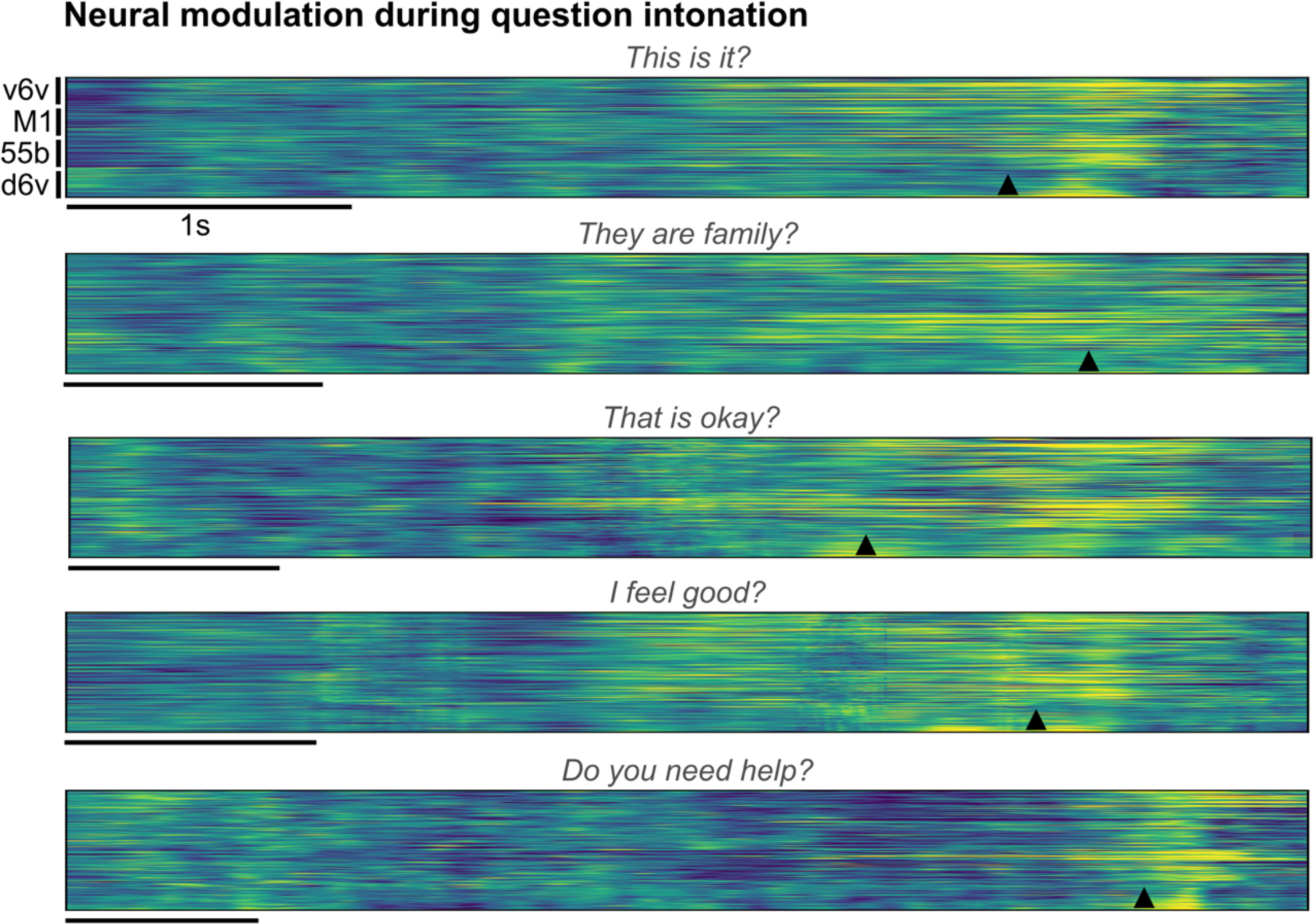
Neural modulation during question intonation. Trial-averaged normalized spike-band power (each row in a group is one electrode) during trials where the participant modulated his intonation to say the cued sentence as a question. Trials with the same cue sentence (n=16) were aligned using dynamic time warping and the mean activity across trials spoken as statements was subtracted to better show the increased neural activity around the intonation-modulation at the end of the sentence. The onset of the word that was pitch-modulated in closed-loop is indicated by the arrowhead at the bottom of each example.

**Extended Data Fig. 7:**
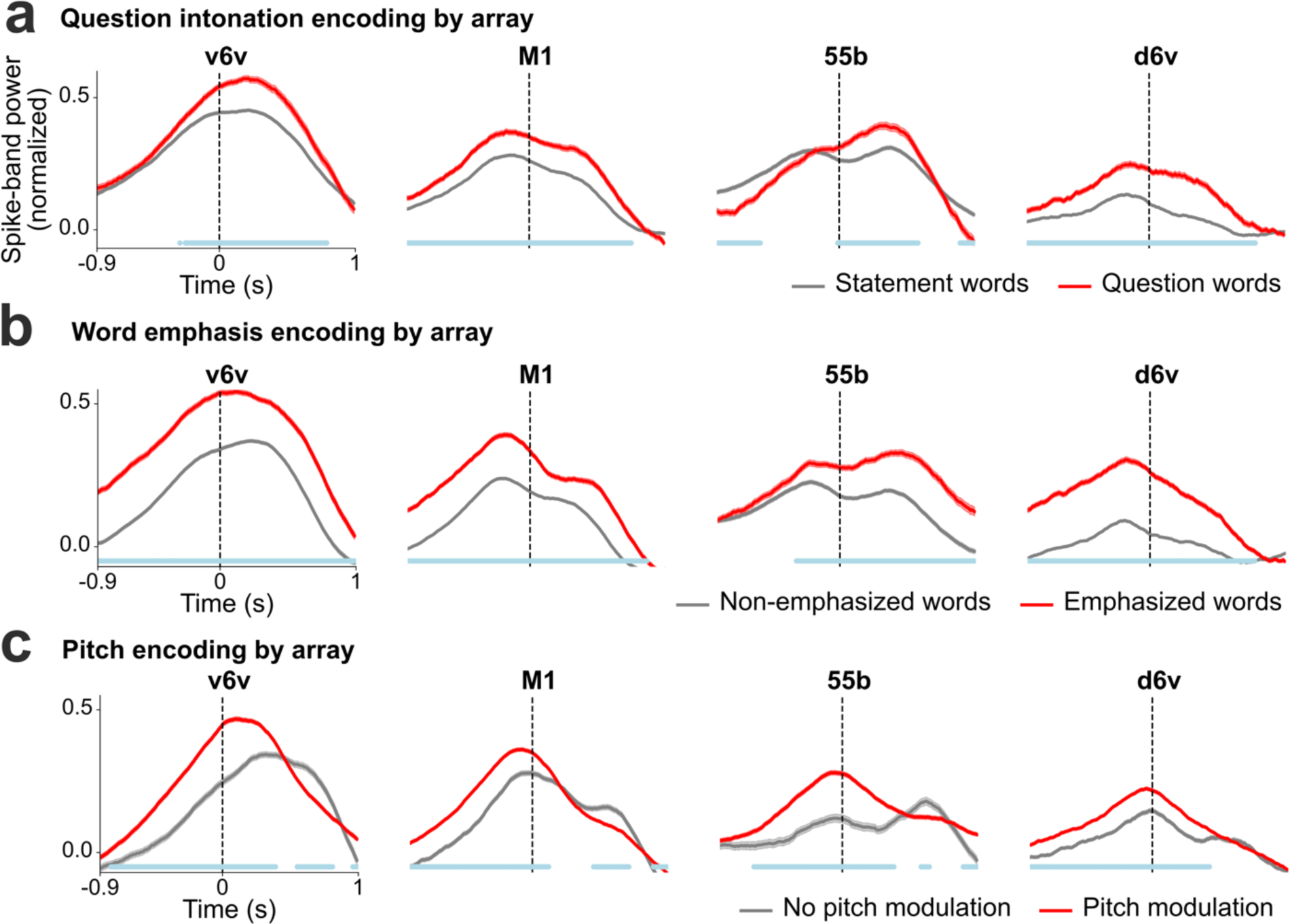
Paralinguistic features encoding recorded from individual arrays. **a.** Trial-averaged spike-band power (mean ± s.e.), averaged across all electrodes within each array, for words spoken as statements and as questions. At every time point, the spike-band power for statement words and question words were compared using the Wilcoxon rank-sum test. The blue line at the bottom indicates the time points where the spike-band power in statement words and question words were significantly different (*p*<0.001, n_1_=970, n_2_=184). **b.** Trial averaged spike-band power across each array for non-emphasized and emphasized words. The spike-band power was significantly different between non-emphasized words and emphasized words at time points shown in blue (*p*<0.001, n_1_=1269, n_2_=333). **c.** Trial-averaged spike-band power across each array for words without pitch modulation and words with pitch modulation (from the three-pitch melodies singing task). Words with low and high pitch targets are grouped together as the ‘pitch modulation’ category (we excluded medium pitch target words where the participant used his normal pitch). The spike-band power was significantly different between no pitch modulation and pitch modulation at time points shown in blue (*p*<0.001, n_1_=486, n_2_=916).

**Extended Data Fig. 8:**
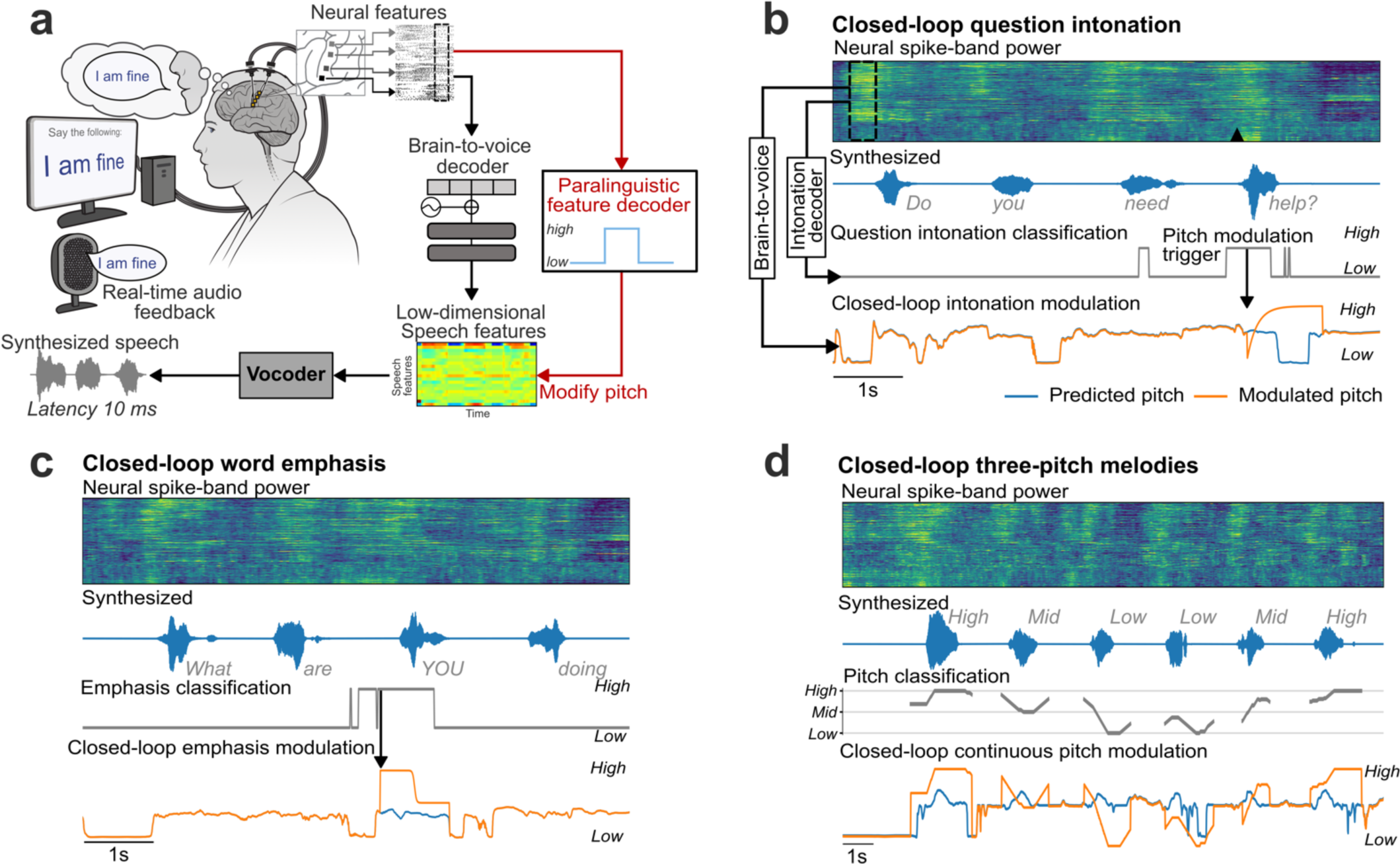
Closed-loop paralinguistic features modulation. **a.** An overview of the paralinguistic feature decoder and pitch modulation pipeline. An independent paralinguistic feature decoder ran in parallel to the regular brain-to-voice decoder. Its output causally modulated the pitch feature predicted by brain-to-voice, resulting in a pitch-modulated voice. **b.** An example trial of closed-loop intonation modulation for speaking a sentence as a question. A separate binary decoder identified the change in intonation and sent a trigger (downward arrow) to modulate the pitch feature output of the regular brain-to-voice decoder according to a predefined pitch profile for asking a question (low pitch to high pitch). Neural activity of an example trial with its synthesized voice output is shown along with the intonation decoder output, time of modulation trigger (downward arrow), originally predicted pitch feature and the modulated pitch feature used for voice synthesis. **c**. An example trial of closed-loop word emphasis where the word “*YOU*” from “*What are YOU doing*” was emphasized. To emphasize a word, we applied a predefined pitch profile (high pitch to low pitch) along with a 20% increase in the loudness of the predicted speech samples. **d**. An example trial of closed-loop pitch modulation for singing a melody with three pitch levels. The three-pitch classifier output was used to continuously modulate the predicted pitch feature output from the brain-to-voice decoder.

**Extended Data Fig. 9:**
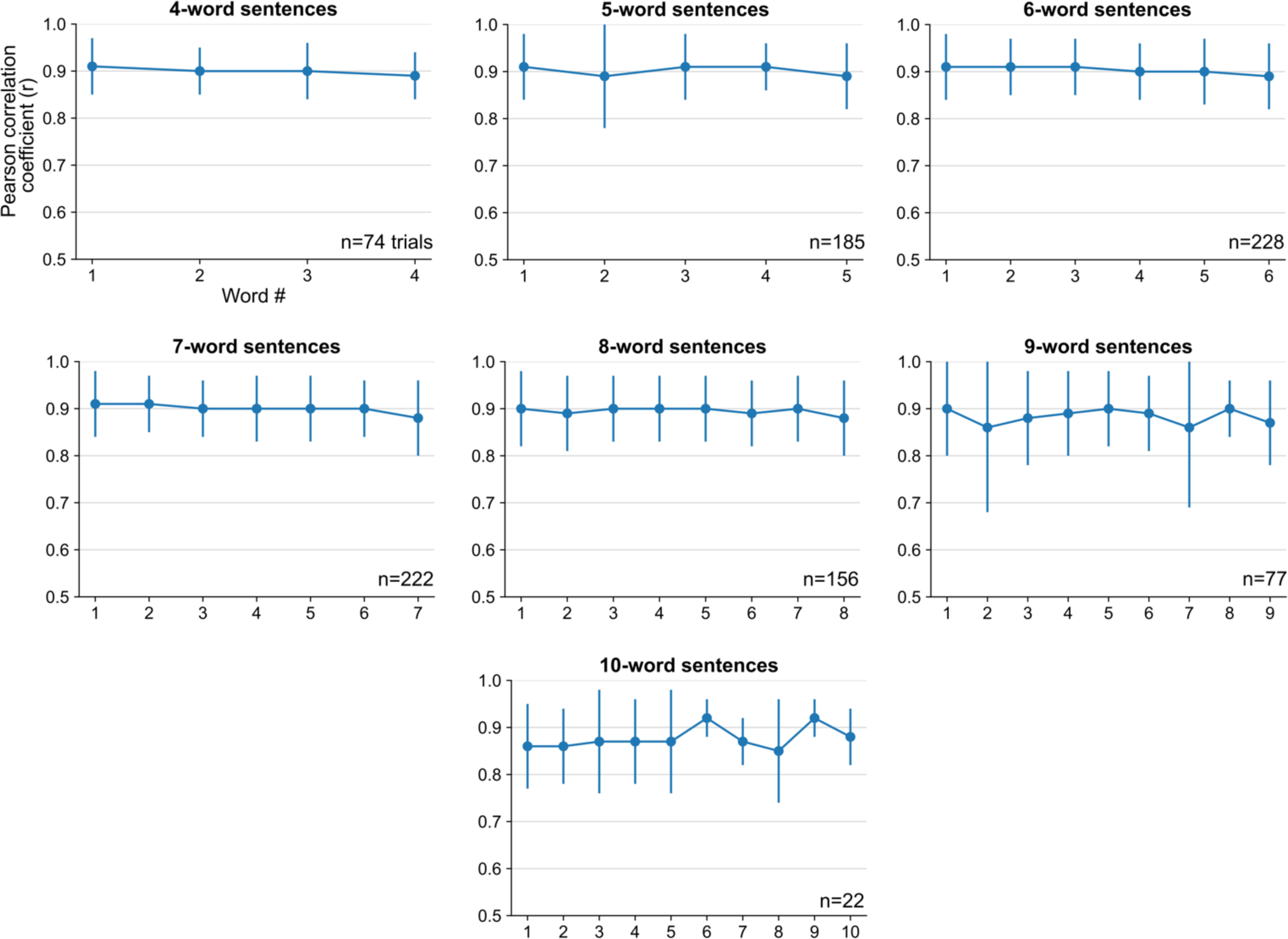
Pearson correlation coefficients over the course of a sentence. Pearson correlation coefficient (*r*) of individual words in sentences of different lengths (mean ± s.d.). The correlation between target and synthesized speech remained consistent throughout the length of sentence, indicating that the quality of synthesized voice was consistent throughout the sentence. Note that there were fewer longer evaluation sentences.

**Extended Data Fig. 10:**
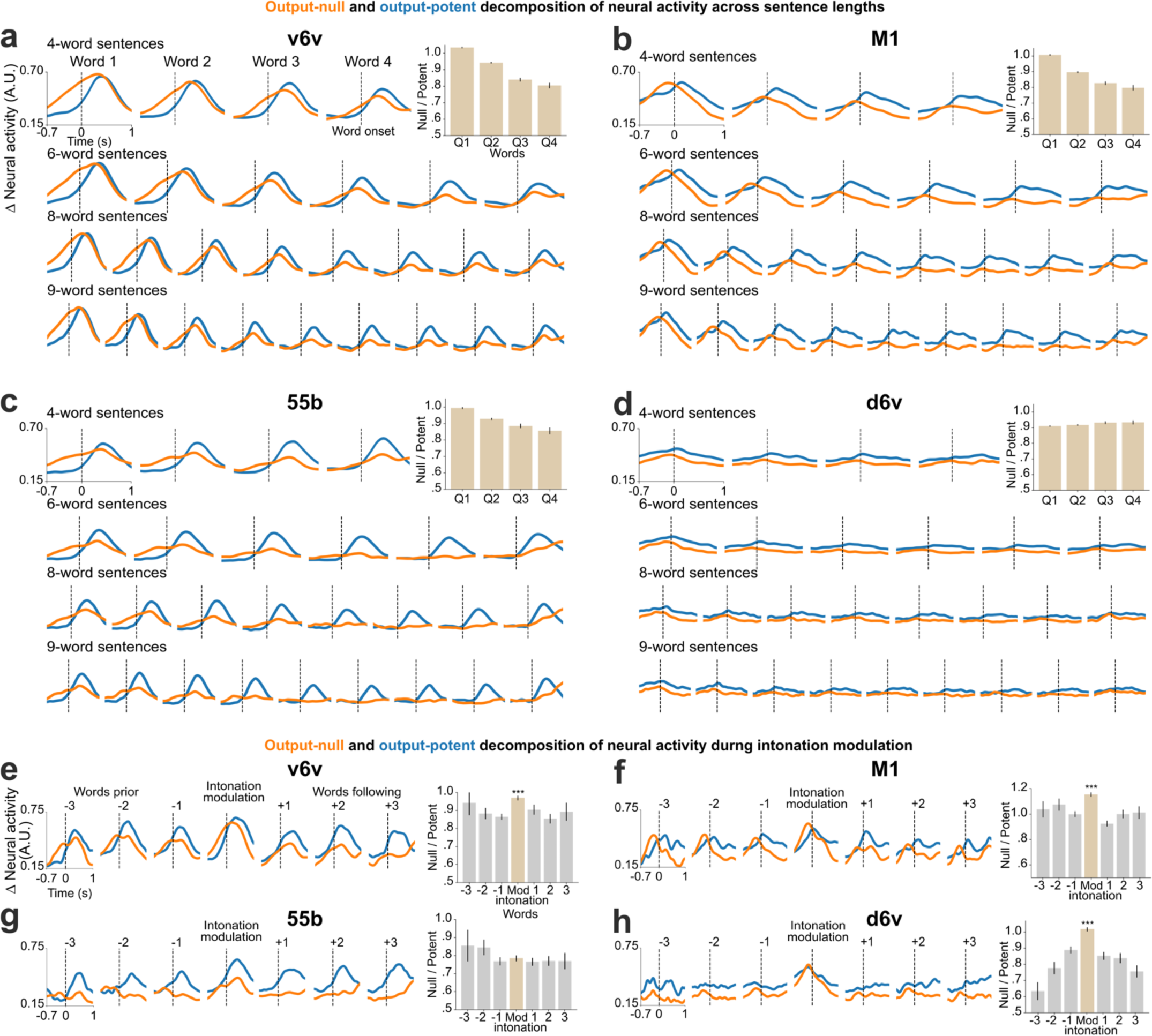
Output-null and output-potent neural dynamics during speech production in individual arrays. a-d. Average approximated output-null (orange) and output-potent (blue) components of neural activity during attempted speech of cued sentences of different lengths. Here the neural components are computed for each array independently by training separate linear decoders (i.e., repeating the analyses of Fig. 4 for individual arrays independently). A subset of sentence lengths are shown in the interest of space. Note that the d6v array had much less speech-related modulation. Bar plots within each panel show a summary of all the data (including the not-shown sentence lengths) by taking the average null/potent activity ratios for words in the first-quarter, second-quarter, third-quarter, and fourth-quarter of each sentence (mean ± s.e.). **e-h.** Average output-null and output-potent activity during intonation modulation (question-asking or word emphasis) computed separately for each array. Output-null activity shows an increase during intonation modulated word in all arrays. Null/potent activity ratios are summarized in bar plots of intonation-modulated word (red) and the words preceding or following it (gray) (mean ± s.e.). The null/potent ratios of modulated words were significantly different from that of non-modulated words for the v6v, M1 and d6v arrays (Wilcoxon rank-sum, v6v: *p*= 10^-11^, M1: *p*= 10^-16^, 55b: *p*= 0.3, d6v: *p*= 10^-26^, n_1_=460, n_2_=922).

## Supplementary media

**Video 1: Dysarthric speech of the participant.** This video shows the participant, who has severe dysarthria due to ALS, attempting to speak the sentences cued on the screen. The participant’s speech is unintelligible to naïve listeners. From post-implant day 27.

Link to view online: https://ucdavis.box.com/s/p8197z804du92225ff0o06l263tjhxcb

**Video 2: Closed-loop voice synthesis during attempted vocalized speech.** This video shows 13 consecutive closed-loop trials of instantaneous voice synthesis as the participant attempted to speak cued sentences. The synthesized voice was played back continuously in real-time through a speaker. From post-implant day 179.

Link to view online: https://ucdavis.box.com/s/esulu85i7meojqnpphr65ioq9pui9xbq

**Video 3: Closed-loop voice synthesis with simultaneous brain-to-text decoding.** This video shows 15 consecutive closed-loop trials of instantaneous voice synthesis with simultaneous brain-to-text decoding that acted as closed-captioning when the participant attempted to speak cued sentences. From post-implant day 110.

Link to view online: https://ucdavis.box.com/s/mw6h5pzvvy6d9kzrt7ayzg1wiag7nxrv

**Video 4: Closed-loop voice synthesis during attempted mimed speech.** This video shows 10 consecutive closed-loop trials of instantaneous voice synthesis with audio feedback as the participant “mimed” the cued sentences without vocalizing. The decoder was not trained on any mimed speech neural data. From post-implant day 195.

Link to view online: https://ucdavis.box.com/s/wpvbw5wogy5kvoalknomfgmf56kxnxmm

**Video 5: Closed-loop voice synthesis during self-initiated free responses.** This video shows 9 closed-loop trials of instantaneous voice synthesis with audio feedback as the participant responded to open-ended questions or was asked to say whatever he wanted. We used this opportunity to ask the participant for his feedback on this brain-to-voice neuroprosthesis. A brain-to-text decoder was used simultaneously to help with understanding what the participant was saying. From post-implant days 172, 179, 186, 188, 193 and 195.

Link to view online: https://ucdavis.box.com/s/bl24hf5kojnz5lm7b6rfq6ejxcm82f0o

**Video 6: Closed-loop own-voice synthesis during attempted speech.** This video shows 9 consecutive closed-loop trials of instantaneous speech synthesis in a voice that sounds like the participant’s own pre-ALS voice as he attempted to speak cued sentences. From post-implant day 286.

Link to view online: https://ucdavis.box.com/s/0vbppq1bevhhblrdfs465fdwuvcn06nd

**Video 7: Closed-loop voice synthesis of pseudo-words.** This video shows 5 consecutive trials of closed-loop synthesis of made-up pseudo-words using the brain-to-voice decoder. The decoder was not trained on any pseudo-words. From post-implant day 179.

Link to view online: https://ucdavis.box.com/s/4qhzyvr0i364xsvaej8zicaf5na4zr44

**Video 8: Closed-loop voice synthesis of interjections.** This video shows 5 trials of closed-loop synthesis of interjections using the brain-to-voice decoder. The decoder was not trained on these words. From post-implant day 186.

Link to view online: https://ucdavis.box.com/s/m234b9ilqpmmcv1yyrmqvchq1z8k3ttl

**Video 9: Closed-loop voice synthesis for spelling words.** This video shows 7 trials of closed-loop synthesis where the participant was spelling cued words one letter at a time using the brain-to-voice decoder. The decoder was not trained on this task. From post-implant day 186.

Link to view online: https://ucdavis.box.com/s/cv8l2ef2t5u4i2km4lwi122z67mxckie

**Video 10: Closed-loop question intonation.** This video shows 10 selected trials where the participant modulated his intonation to say a sentence as a question (indicated by ‘?’ in the cue) or as a statement by using an intonation decoder that modulated the brain-to-voice synthesis in closed-loop. From post-implant day 286.

Link to view online: https://ucdavis.box.com/s/67xnduo76355v93nzjuvez5utll5qung

**Video 11: Closed-loop word emphasis.** This video shows 8 selected trials where certain (capitalized) words in the cued sentences were emphasized by the participant by using an emphasis decoder that modulated the brain-to-voice synthesis in closed-loop. From post-implant day 286.

Link to view online: https://ucdavis.box.com/s/s7crvym9q9dro5mo9a6wmlltjuy0c88f

**Video 12: Singing three-pitch melodies in closed-loop.** This video shows 3 consecutive trials where the participant sung short melodies with three pitch targets by using a pitch decoder that modulated the brain-to-voice synthesis in closed-loop. At the start of each trial, an audio cue plays the target melody. The on-screen targets then turn from red to green to indicate that the participant should begin. The vertical bar on the left of the screen shows the instantaneous decoded pitch (low, mid, high). Additionally, interactive visual cues for each pitch target are shown on the screen. These visual feedback cues show the note in the melody that the participant is singing. From post-implant day 342.

Link to view online: https://ucdavis.box.com/s/quj4z50adoibkfysgse21b6t5jzk7xmp

**Video 13: Singing three-pitch melodies in closed-loop using a unified brain-to-voice decoder.** This video shows 3 trials where the participant sung short melodies with three pitch targets by using a single unified brain-to-voice decoder that inherently synthesizes intended pitch in closed-loop. At the start of each trial, an audio cue plays the target melody. The vertical bar on the left of the screen shows the instantaneous decoded pitch (low, mid, high) for visual feedback only (i.e., this separately-decoded pitch, which is the same as in Video 12, is not used in the unified brain-to-voice model). Interactive visual cues show the note in the melody that the participant is singing, providing visual feedback. From post-implant day 342.

Link to view online: https://ucdavis.box.com/s/qu5nwz8qg6hpxtqnvjqkxla1mhoic99c

**Video 14: Closed-loop voice synthesis in session 1.** This video shows 3 closed-loop trials of instantaneous voice synthesis from the participant’s first day of neural recording (post-implant day 25). The brain-to-voice decoder was trained during this session using 190 sentence trials from a limited 50-word vocabulary recorded earlier on the same day. The second part of the video shows the same three trials reconstructed offline using an optimized brain-to-voice decoder (i.e., the algorithm used throughout the rest of this manuscript), which improved intelligibility.

Link to view online: https://ucdavis.box.com/s/aw59fr2kddxkyagw7phmobg1d1hiwp9d

**Audio 1: Acausal speech synthesis by predicting discrete speech units.** This audio shows 3 example trials of speech reconstructed offline using the approach of predicting discrete speech units acausally at the end of the sentence using CTC loss. From post-implant day 25.

Link to view online: https://ucdavis.box.com/s/b0r5n00n0rss0fjdk4b1xuzf1gvy3mwn

## Notes

### Summary of Updates

Two additional extended data figures are included.

## References

1. Card, N. S. et al. An Accurate and Rapidly Calibrating Speech Neuroprosthesis. N. Engl. J. Med. 391, 609–618 (2024).

2. Willett, F. R. et al. A high-performance speech neuroprosthesis. Nature 620, 1031–1036 (2023).

3. Metzger, S. L. et al. A high-performance neuroprosthesis for speech decoding and avatar control. Nature 620, 1037– 1046 (2023).

4. Silva, A. B., Littlejohn, K. T., Liu, J. R., Moses, D. A. & Chang, E. F. The speech neuroprosthesis. Nat. Rev. Neurosci. 25, 473–492 (2024).

5. Herff, C. et al. Generating Natural, Intelligible Speech From Brain Activity in Motor, Premotor, and Inferior Frontal Cortices. Front. Neurosci. 13, (2019).

6. Angrick, M. et al. Speech synthesis from ECoG using densely connected 3D convolutional neural networks. J. Neural Eng. 16, 036019 (2019).

7. Anumanchipalli, G. K., Chartier, J. & Chang, E. F. Speech synthesis from neural decoding of spoken sentences. Nature 568, 493–498 (2019).

8. Meng, K. et al. Continuous synthesis of artificial speech sounds from human cortical surface recordings during silent speech production. J. Neural Eng. 20, 046019 (2023).

9. Le Godais, G. et al. Overt speech decoding from cortical activity: a comparison of different linear methods. (2023) doi:10.3389/fnhum.2023.1124065.

10. Liu, Y. et al. Decoding and synthesizing tonal language speech from brain activity. Sci. Adv. 9, eadh0478 (2023).

11. Berezutskaya, J. et al. Direct speech reconstruction from sensorimotor brain activity with optimized deep learning models. J. Neural Eng. 20, 056010 (2023).

12. Chen, X. et al. A neural speech decoding framework leveraging deep learning and speech synthesis. *Nat*. Mach. Intell. 6, 467–480 (2024).

13. Wilson, G. H. et al. Decoding spoken English from intracortical electrode arrays in dorsal precentral gyrus. J. Neural Eng. 17, 066007 (2020).

14. Wairagkar, M., Hochberg, L. R., Brandman, D. M. & Stavisky, S. D. Synthesizing Speech by Decoding Intracortical Neural Activity from Dorsal Motor Cortex. in 2023 11th International IEEE/EMBS Conference on Neural Engineering (NER) 1–4 (2023). doi:10.1109/NER52421.2023.10123880.

15. Angrick, M. et al. Real-time synthesis of imagined speech processes from minimally invasive recordings of neural activity. *Commun*. Biol. 4, 1–10 (2021).

16. Wu, X., Wellington, S., Fu, Z. & Zhang, D. Speech decoding from stereo-electroencephalography (sEEG) signals using advanced deep learning methods. J. Neural Eng. 21, 036055 (2024).

17. Angrick, M. et al. Online speech synthesis using a chronically implanted brain–computer interface in an individual with ALS. Sci. Rep. 14, 9617 (2024).

18. Glasser, M. F. et al. A multi-modal parcellation of human cerebral cortex. Nature 536, 171–178 (2016).

19. Vaswani, A., et al. Attention Is All You Need. Preprint at 10.48550/arXiv.1706.03762 (2023).

20. Downey, J. E., Schwed, N., Chase, S. M., Schwartz, A. B. & Collinger, J. L. Intracortical recording stability in human brain–computer interface users. J. Neural Eng. 15, 046016 (2018).

21. Valin, J.-M. & Skoglund, J. LPCNET: Improving Neural Speech Synthesis through Linear Prediction. in ICASSP 2019 - 2019 IEEE International Conference on Acoustics, Speech and Signal Processing (ICASSP) 5891–5895 (2019). doi:10.1109/ICASSP.2019.8682804.

22. Li, Y. A., Han, C., Raghavan, V. S., Mischler, G. & Mesgarani, N. StyleTTS 2: Towards Human-Level Text-to-Speech through Style Diffusion and Adversarial Training with Large Speech Language Models. Preprint at 10.48550/arXiv.2306.07691 (2023).

23. Dichter, B. K., Breshears, J. D., Leonard, M. K. & Chang, E. F. The Control of Vocal Pitch in Human Laryngeal Motor Cortex. Cell 174, 21–31.e9 (2018).

24. Kaufman, M. T., Churchland, M. M., Ryu, S. I. & Shenoy, K. V. Cortical activity in the null space: permitting preparation without movement. Nat. Neurosci. 17, 440–448 (2014).

25. Stavisky, S. D., Kao, J. C., Ryu, S. I. & Shenoy, K. V. Motor Cortical Visuomotor Feedback Activity Is Initially Isolated from Downstream Targets in Output-Null Neural State Space Dimensions. Neuron 95, 195–208.e9 (2017).

26. Churchland, M. M. & Shenoy, K. V. Preparatory activity and the expansive null-space. Nat. Rev. Neurosci. 25, 213– 236 (2024).

27. Moses, D. A. et al. Neuroprosthesis for Decoding Speech in a Paralyzed Person with Anarthria. N. Engl. J. Med. 385, 217–227 (2021).

28. Bouchard, K. E., Mesgarani, N., Johnson, K. & Chang, E. F. Functional organization of human sensorimotor cortex for speech articulation. Nature 495, 327–332 (2013).

29. Chartier, J., Anumanchipalli, G. K., Johnson, K. & Chang, E. F. Encoding of Articulatory Kinematic Trajectories in Human Speech Sensorimotor Cortex. Neuron 98, 1042–1054.e4 (2018).

30. Lu, J. et al. Neural control of lexical tone production in human laryngeal motor cortex. Nat. Commun. 14, 6917 (2023).

31. Breshears, J. D., Molinaro, A. M. & Chang, E. F. A probabilistic map of the human ventral sensorimotor cortex using electrical stimulation. (2015) doi:10.3171/2014.11.JNS14889.

32. Ammanuel, S. G. et al. Intraoperative cortical stimulation mapping with laryngeal electromyography for the localization of human laryngeal motor cortex. (2024) doi:10.3171/2023.10.JNS231023.

33. Pandarinath, C. et al. Neural population dynamics in human motor cortex during movements in people with ALS. eLife 4, e07436 (2015).

34. Stavisky, S. D. et al. Neural ensemble dynamics in dorsal motor cortex during speech in people with paralysis. eLife 8, e46015 (2019).

35. Willett, F. R. et al. Hand Knob Area of Premotor Cortex Represents the Whole Body in a Compositional Way. Cell 181, 396–409.e26 (2020).

## References

36. Ali, Y. H., et al. BRAND: a platform for closed-loop experiments with deep network models. J. Neural Eng. 21, 026046 (2024).

37. Young, D. et al. Signal processing methods for reducing artifacts in microelectrode brain recordings caused by functional electrical stimulation. J. Neural Eng. 15, 026014 (2018).

38. Levelt, W. J., Roelofs, A. & Meyer, A. S. A theory of lexical access in speech production. Behav. Brain Sci. 22, 1–38; discussion 38-75 (1999).

39. Räsänen, O., Doyle, G. & Frank, M. C. Unsupervised word discovery from speech using automatic segmentation into syllable-like units. in 3204–3208 (2015). doi:10.21437/Interspeech.2015-645.

40. Williams, A. H. et al. Discovering Precise Temporal Patterns in Large-Scale Neural Recordings through Robust and Interpretable Time Warping. Neuron 105, 246–259.e8 (2020).

